# Representational drift in the mouse visual cortex

**DOI:** 10.1101/2020.10.05.327049

**Authors:** Daniel Deitch, Alon Rubin, Yaniv Ziv

**Author notes:** Corresponding Author: Yaniv Ziv.

## Abstract

Neuronal representations in the hippocampus and related structures gradually change over time despite no changes in the environment or behavior. The extent to which such ‘representational drift’ occurs in sensory cortical areas and whether the hierarchy of information flow across areas affects neural-code stability have remained elusive. Here, we address these questions by analyzing large-scale optical and electrophysiological recordings from six visual cortical areas in behaving mice that were repeatedly presented with the same natural movies. We found representational drift over timescales spanning minutes to days across multiple visual areas. The drift was driven mostly by changes in individual cells’ activity rates, while their tuning changed to a lesser extent. Despite these changes, the structure of relationships between the population activity patterns remained stable and stereotypic, allowing robust maintenance of information over time. Such population-level organization may underlie stable visual perception in the face of continuous changes in neuronal responses.

## Introduction

One of the great marvels of the brain is that it achieves persistent functionality throughout adult life despite an extensive continuous turnover of its bio-molecular and cellular building blocks (Yasumatsu *et al*., 2008; Holtmaat and Svoboda, 2009; Minerbi *et al*., 2009; Loewenstein, Kuras and Rumpel, 2011; Alvarez-Castelao and Schuman, 2015). Recent advances in electrophysiology and optical imaging techniques enable to study in awake behaving animals the persistence over time of neuronal coding properties, such as the tuning of neurons to specific stimuli (Rokni *et al*., 2007; Tolias *et al*., 2007; Bondar *et al*., 2009; Andermann, Kerlin and Reid, 2010; Huber *et al*., 2012; Ziv *et al*., 2013; Peron *et al*., 2015; Poort *et al*., 2015; Okun *et al*., 2016; Dhawale *et al*., 2017; Jun *et al*., 2017). Some of these studies exposed a substantial degree of variability in neuronal responses to the same stimuli over timescales spanning minutes to weeks, prompting neuroscientists to question the naïve assumption that stable neuronal codes are essential for stable brain functionality (Tolhurst, Movshon and Dean, 1983; Arieli *et al*., 1996; Rokni *et al*., 2007; Faisal, Selen and Wolpert, 2008; Minerbi *et al*., 2009; Cohen and Maunsell, 2010; Huber *et al*., 2012; Ziv *et al*., 2013; Lütcke, Margolis and Helmchen, 2013; Montijn, Goltstein and Pennartz, 2015; Rubin *et al*., 2015; Schölvinck *et al*., 2015; Rose *et al*., 2016; Chambers and Rumpel, 2017; Clopath *et al*., 2017; Dhawale *et al*., 2017; Driscoll *et al*., 2017; Engel and Steinmetz, 2019; Rule *et al*., 2020; Sheintuch *et al*., 2020).

One example is the neuronal representations of space in the hippocampus, which gradually change over time despite no apparent changes in the spatial environment or behavior (Ziv *et al*., 2013; Mankin *et al*., 2015; Rubin *et al*., 2015; Sheintuch *et al*., 2020). Similar continuous dynamics of neuronal representations has also been shown in other brain structures (Driscoll *et al*., 2017). The finding of this so called ‘representational drift’ (Rule, O’Leary and Harvey, 2019) was surprising, because classical models of memory consider the stability of the engram as the basis for the persistence of memory (Josselyn, Köhler and Frankland, 2015; Tonegawa, Morrissey and Kitamura, 2018). Notably, representational drift is different than mere variability in neuronal responses in that the changes in the cells’ coding properties are gradual. Namely, in representational drift, the similarity between any two representations of the same stimulus gradually decays with elapsed time; in contrast, variability in neuronal responsiveness does not lead to such time-dependent decay in the similarity between representations (Clopath *et al*., 2017; Rule, O’Leary and Harvey, 2019).

In the hippocampus, representational drift is driven primarily by changes in the activity rates of individual neurons, whereas the cells’ tuning to specific positions changes to a lesser extent (Ziv *et al*., 2013; Rubin *et al*., 2015). These findings suggest that coding stability is a complex trait that is affected by different aspects of cellular physiology and connectivity. The specific mechanisms that underlie representational drift remain elusive, but it was suggested that drift may be an inevitable outcome of the network dynamics in deep brain circuits that consist of multiple input and output loops (Rule, O’Leary and Harvey, 2019). Consistent with this logic, and given the need to support stable perception and motor outputs, it is plausible that brain circuits situated closer to the sensory input or to the motor output will display highly stable neuronal representations (Haak, Morland and Engel, 2015; Haak and Mesik, 2016).

While a direct examination of this hypothesis is still lacking, several recent chronic two-photon imaging studies of sensory cortices investigated coding stability and found variability in neuronal responsiveness to sensory stimuli over multiple days (Andermann, Kerlin and Reid, 2010; Montijn *et al*., 2016; Rose *et al*., 2016; Ranson, 2017; Jeon *et al*., 2018). For instance, in the primary visual cortex (V1), Rose et al. (2016) revealed session-to-session, time-independent variability of neuronal visual tuning properties (e.g., ocular dominance). Similarly, Montijn et al. (2016) reported that the neuronal responses in V1 are variable across trials within the same day, but their tuning is relatively stable across days.

These studies provide clear indications that representations of visual stimuli in L2/3 neurons of V1 are variable over time. However, it remains unclear if and to what extent the visual cortex (or any sensory cortex, for that matter) exhibits representational drift that is similar to that observed in deep circuits (Ziv *et al*., 2013; Rubin *et al*., 2015), in terms of the degree to which different aspects of cells’ coding properties, such as tuning and activity rate, change over time. It is also unknown whether higher-order sensory cortical areas generate representations that are less stable than those of primary sensory cortices, and how the stability of neuronal coding properties differs across different layers within a given cortical area.

Recently, the Allen Brain Institute published two large-scale, standardized physiological surveys of neuronal coding in the visual cortex (Allen Brain Observatory) (Siegle *et al*., 2019; de Vries *et al*., 2020). These datasets consist of optical and electrophysiological recordings of tens of thousands of neurons from six different visual cortical areas in hundreds of awake behaving mice that were repeatedly presented with the same set of stimuli. Thus, they offer a unique opportunity to study coding stability across different areas of the visual cortex and over different timescales, from minutes to days. The dense recording using Neuropixels probes (Jun *et al*., 2017) allows within-mouse comparison across visual areas, whereas the two-photon Ca^2+^ imaging dataset enables the longitudinal analysis of a large population of the same cells, both within and across days. Given that the study of coding stability is often confounded by technical issues, such as the mechanical stability of the recoding apparatus, the fact that the same experiments were conducted using two different recording techniques (Neuropixels and two-photon Ca^2+^ imaging) can help overcome and control for limitations and biases associated with each technique. Furthermore, a specific set of stimuli – natural scene movies – were used in these experiments and on different days. This allows the detailed investigation of the stability of visual representations that are more complex and ethologically relevant than the synthetic stimuli traditionally used for longitudinal studies (David, Vinje and Gallant, 2004; Kampa *et al*., 2011; Talebi and Baker, 2012).

Here, we utilized these datasets to address basic questions regarding the stability and dynamics of visual representations. We found that representational drift occurs across different visual areas, over timescales spanning minutes to days, and is primarily driven by changes in the cells’ activity rates rather than in their tuning. We demonstrate that despite clear time-dependent changes in neuronal responsiveness to visual stimuli, the structure of relationships between neuronal population activity patterns remained stable, permitting the conservation of visual information over time.

## Results

We analyzed publicly available datasets by the Allen Brain Observatory from experiments that used two recording techniques: two-photon Ca^2+^ imaging (de Vries *et al*., 2020) and electrophysiology via Neuropixels probes (Siegle *et al*., 2019). The Ca^2+^ imaging dataset comprises neuronal activity from nearly 60,000 neurons collected from six visual cortical areas, across different layers, from hundreds of adult mice that were presented with the same set of visual stimuli (Fig. 1A-D). Each mouse was imaged from a single cortical area while performing three imaging sessions, separated by a different number of days. During each imaging session, mice viewed a battery of natural and artificial stimuli (Fig. 1C and Methods). The Neuropixels dataset comprises neuronal activity from nearly 90,000 single units collected from six visual areas, thalamic nuclei, and the hippocampus, from 58 adult mice (Fig. 1E-H). Each mouse was implanted with multiple Neuropixels probes in different brain areas and went through a single recording session while viewing a battery of natural and artificial stimuli (Fig. 1E-F). In this study, we focused our analysis on neuronal activity recorded during the presentations of two natural movies, because they were presented in all imaging sessions across days (‘Natural Movie 1’ in the Ca^2+^ imaging dataset) or twice within the same recording session (‘Natural Movie 3’ in the Ca^2+^ imaging and Neuropixels datasets; and ‘Natural Movie 1’ in the Neuropixels dataset). This experimental design enabled us to study the stability of neuronal representations on different time scales: (1) Between movie repetitions within a single block across seconds-minutes; (2) Between different blocks within the same recording session across minutes-hours; And (3) across sessions recorded on different days. In datasets from both recording techniques, we could readily identify neurons that displayed reliable and distinct tuning curves that were stable across different movies repeats, blocks and days (Fig. 1D, H).

**Figure 1.**
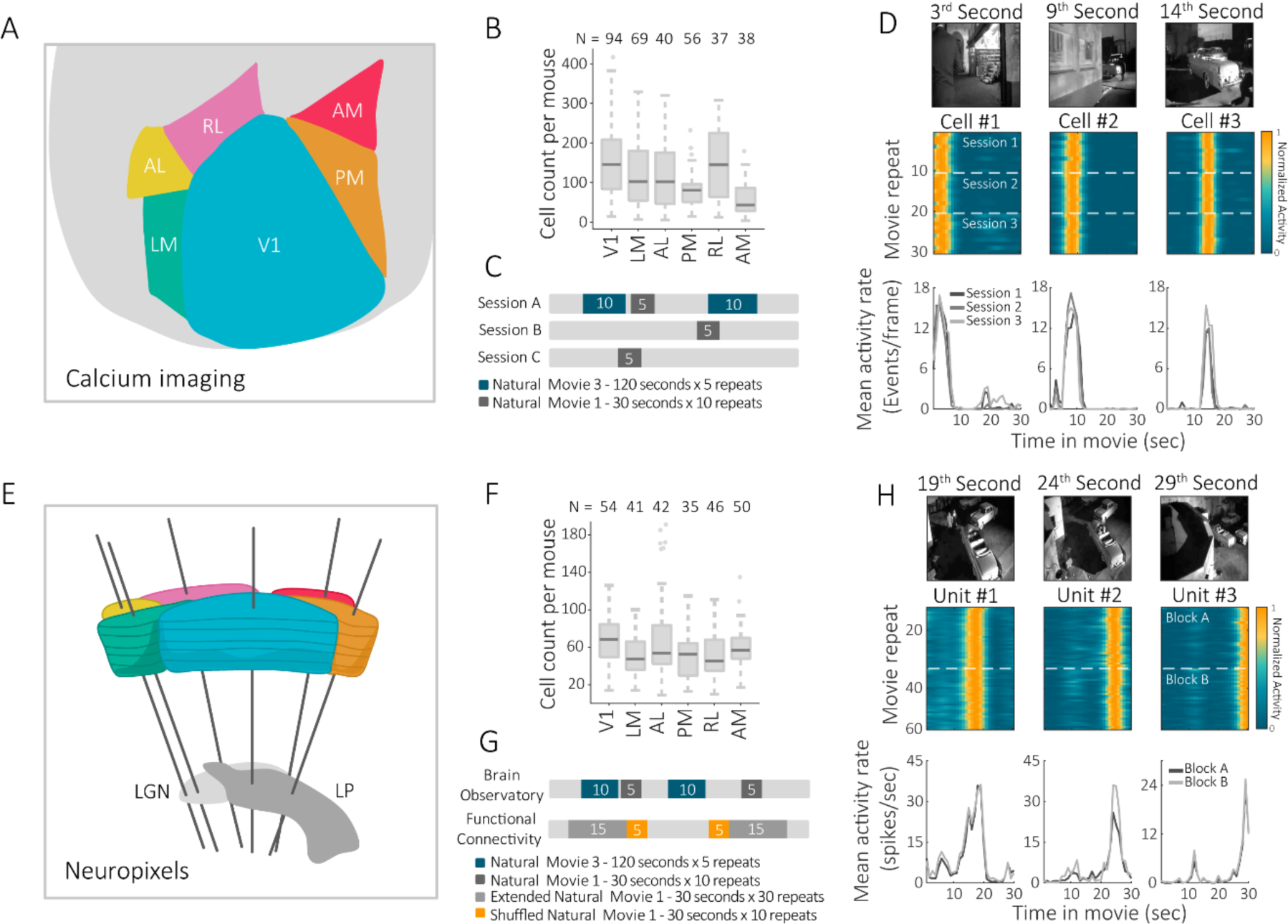
Neurons recorded from various visual cortical areas show reliable tuning to natural movies. **A-D**: Ca^2+^ imaging dataset. **A**. Schematic illustration of the different brain areas imaged using two-photon Ca^2+^ imaging. V1 - Primary visual area, LM – lateral-medial visual area, AL - anterolateral visual area, PM - posteromedial visual area, RL - rostrolateral visual area, AM - anteromedial visual area. **B**. Distribution of cell counts per mouse across brain areas for the two-photon Ca^2+^ imaging dataset. **C**. Experimental design of the Ca^2+^imaging dataset. Each mouse performed three sessions in a random order, separated by a different number of days. Only the two indicated stimuli (‘Natural movie 1’ and ‘Natural movie 3’) were used in our analysis (other stimuli are specified in the Methods). **D**. Responsiveness of three example cells across different ‘Natural movie 1’ repeats spanning three Ca^2+^ imaging sessions. **E-H**: Neuropixels dataset. **E**. Schematic illustration (adapted from Siegle et al. (2019)) of the different brain area recordings using Neuropixels probes. LGN – lateral geniculate nucleus. LP – lateral parietal nucleus. **F**. Distribution of cell counts per mouse across brain areas for the Neuropixels dataset. **G**. Experimental design of the Neuropixels dataset. Thirty-two of the mice performed the ‘Brain Observatory’ battery and 26 performed the ‘Functional Connectivity’ battery. Only the indicated stimuli (‘Natural movie 1’, ‘Natural movie 3’, and ‘Shuffled natural movie 1’) were used in our analysis (other stimuli are specified in the Methods). **H**. Responsiveness of three example cells across different ‘Natural movie 1’ repeats spanning two blocks within the same Neuropixels recording session.

To study the stability and dynamics of visual representations over timescales of seconds to minutes, we analyzed data recorded using Neuropixels probes during the presentations of ‘Natural Movie 1’. We divided each movie repeat into equal time bins and constructed population vectors (PVs) of neuronal activity for each time bin (see Methods). We then calculated the correlation across the PVs of all time bins of all movie repeats as a measure for similarity between neuronal representations across time (Fig. 2A). We found higher PV correlations between the same time bins across movies repeats than between different time bins, indicating distinct and stable representation of the movie sequence (Fig. 2A inset). The average PV correlation values across the same time bins capture the stability of the ensemble representation between different movie repeats (Fig. 2B). If the stability of the visual representation is mostly affected by the variability of the neuronal responses, then the PV correlation values across two movie repeats that are close in time would be similar to those of two movies repeats that are more remote in time. If, however, the neuronal representation is drifting, then the PV correlation values across two movie repeats that are close in time would be higher than those of two movies repeats that are more remote in time (Clopath *et al*., 2017). Calculating the mean PV correlation as a function of the interval between movie repeats showed a significant gradual decline, indicating representational drift in all studied visual areas (Fig. 2C-E and Supp. Fig. 1A).

**Figure 2.**
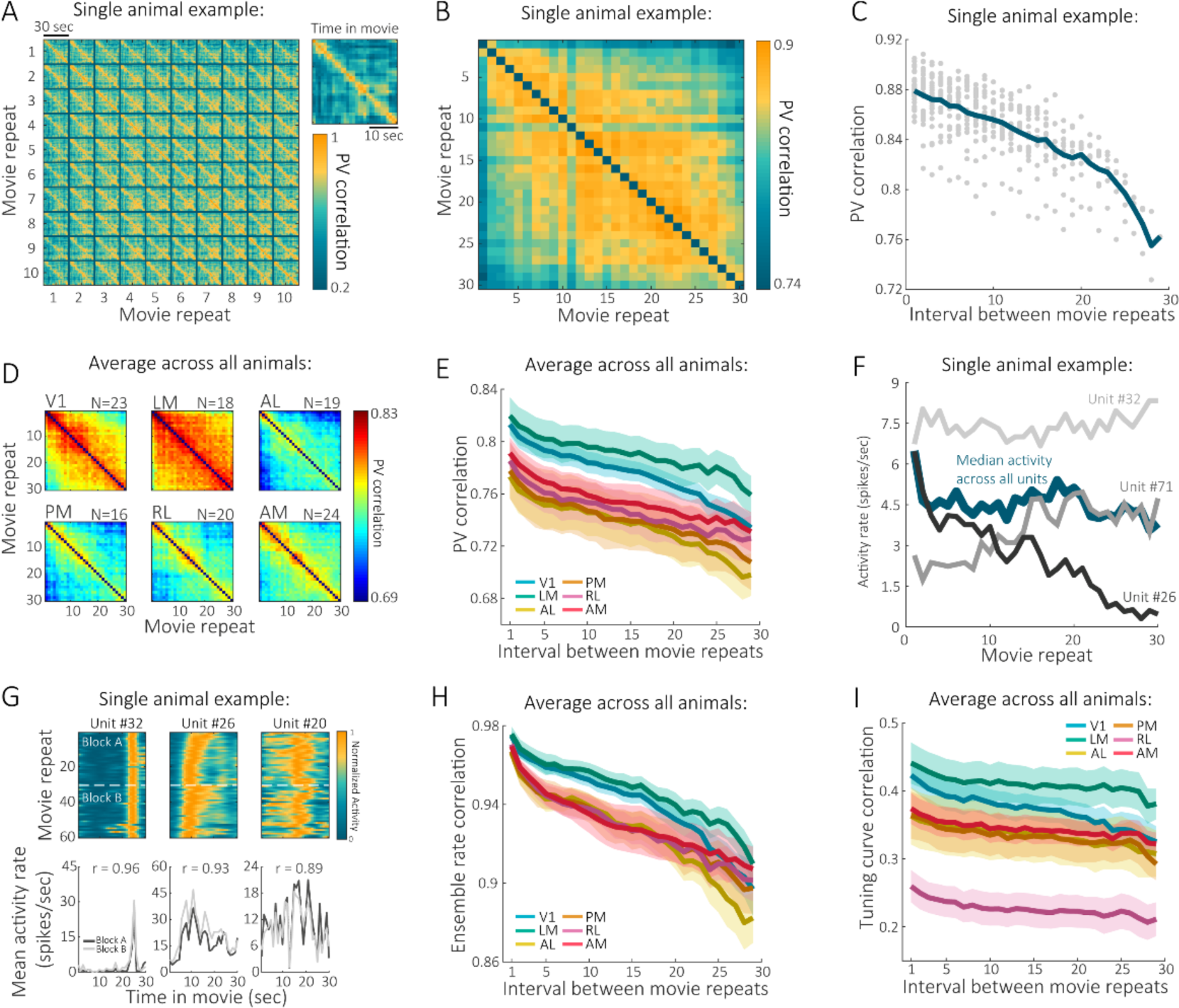
The visual cortex exhibits representational drift across subsequent presentations of a natural stimulus over timescales of seconds-minutes. **A**. Population vector (PV) correlation between the first 10 (out of 30) repeats of ‘Natural movie 1’ of the first block, recorded using a Neuropixels probe, from area PM of a single representative mouse. Inset: the average PV correlation over all pairs across different movie repeats. **B**. Mean PV correlation for each pair of movie repeats from the same mouse shown in A. Each entry is the average PV correlation between all matching time bins for a pair of movie repeats. For visualization, the main diagonal (in which values are equal to 1 by definition) was removed. **C**. Mean PV correlation as a function of the interval between movie repeats. Each data point represents the mean PV correlation value for a single pair of movie repeats from B. **D**. Mean PV correlation between movie repeats across animals and brain areas for the Neuropixels ‘Functional connectivity’ group. **E**. PV correlation as a function of the interval between movie repeats for six visual cortical areas; the difference in PV correlations between the interval of one movie repeat and that of 29 movie repeats was significant for all areas (p < 10^−3^, two-tailed Wilcoxon signed-rank test with Holm–Bonferroni correction). **F**. Mean activity rates for three example units from area PM from the same representative mouse across movie repeats. Each unit exhibited different changes in its activity rates relative to the activity levels of the entire recorded population, which remained relatively stable throughout the block. **G**. Responsiveness of three V1 example cells from the same representative mouse across different repeats of ‘Natural movie 1’, spanning two blocks within the same recording session. Each unit exhibits a different degree of tuning curve stability within a given block and across the two blocks (indicated by the Pearson correlation values in the bottom panels). **H**. Ensemble rate correlation across animals as a function of the interval between movie repeats for six visual cortical areas; the difference in ensemble rate correlations between the interval of one movie repeat and that of 29 movie repeats was significant for all areas (p < 10^−3^, two-tailed Wilcoxon signed-rank test with Holm– Bonferroni correction). **I**. Tuning curve correlation across animals as a function of the interval between movie repeats for six visual cortical areas; the difference in tuning curve correlations between the interval of one movie repeat to that of 29 movie repeats was significant for all areas (p ≤ 0.017, two-tailed Wilcoxon signed-rank test with Holm–Bonferroni correction). Data in panels E, H, I are mean ± SEM across mice.

Could the observed representational drift merely reflect changes in behavioral state or global fluctuation in neuronal activity levels? Indeed, we found a mild drop in running speed and pupil area after the first few movie repeats, potentially reflecting changes in arousal (Niell and Stryker, 2010; Keller, Bonhoeffer and Hübener, 2012; Ayaz *et al*., 2013; Polack, Friedman and Golshani, 2013; Erisken *et al*., 2014; Ruff and Cohen, 2014; Vinck *et al*., 2015; Mineault *et al*., 2016; Dipoppa *et al*., 2018; Musall *et al*., 2019)(Supp. Fig. 2A,B). We also observed a modest decrease in the global neuronal activity rates during the first several movie repeats (Supp. Fig. 2C,D). To test the possibility that these changes in behavior and neuronal activity rates underlie our measurements of representational drift, we removed the first several movie repeats and repeated our analysis. The decline in PV correlation values as a function of the interval between movie repeats in the subsampled dataset was similar to the decline seen in the analysis of the complete dataset (Supp. Fig. 2D,E). Furthermore, the distribution of the differences in activity rates of individual neurons between the beginning and end of each block of movie repeats was centered around zero, suggesting that representational drift is not driven by a systematic decline in firing rates in individual neurons (Supp. Fig. 2F). Thus, changes in the behavioral state or in global neuronal activity levels did not underlie the observed representational drift.

What cellular properties could underlie the observed representational drift? Time-dependent decline in PV correlations may stem from changes in cellular excitability (Fig. 2F) or from changes in the tuning of individual neurons to the presented stimuli (Fig. 2G). To test the contribution of each of these factors to the observed changes in PV correlations over time, we performed two complementary analyses: (1) ‘Ensemble rate correlation’: For each movie repeat we constructed a single vector constituting the overall activity rates of each cell in the recorded population. We then quantified the correlations across all pairs of these vectors, which captured the changes in the cells’ activity rates, irrespective of their tuning to different time points along the movie. This analysis revealed a significant decline in the ensemble rate correlation as a function of the interval between movie repeats in all studied visual areas (Fig. 2H, and Supp. Fig. 1B). (2) ‘Tuning curve correlation’: For each neuron, at each movie repeat, we constructed a vector representing its responsiveness to different time bins of the presented movie (i.e., its tuning curve) and then correlated the tuning curves for the same neurons across different movie repeats. This analysis revealed a modest, yet significant, decline in the tuning curve correlation values as a function of the distance between movie repeats in all studied visual areas (Fig. 2I, and Supp. Fig. 1C). We found similar time-dependent changes in visual tuning when we trained a decoder (*k*-nearest neighbors; see Methods) to infer the time bin associated with a given activity pattern at a given movie repeat based on the activity patterns of the neuronal population during the preceding repeat (Supp. Fig. 3A-F). To minimize the contribution of recording instability to our observations, we restricted our analysis only to cells whose tuning curves were highly correlated across different blocks, thereby ensuring we tracked the same cells within a given block. Even when using this inclusion criterion, we found time-dependent changes in visual representations in all studied cortical areas (Supp. Fig. 4A-F). Notably, we obtained similar results in the Ca^2+^ imaging dataset, further substantiating that the observed drift is not due to recording instability (Supp. Fig. 1D-F). Overall, changes in both the cells’ tuning and activity rates contributed to drift in visual representations over timescales of seconds-minutes.

**Figure 3.**
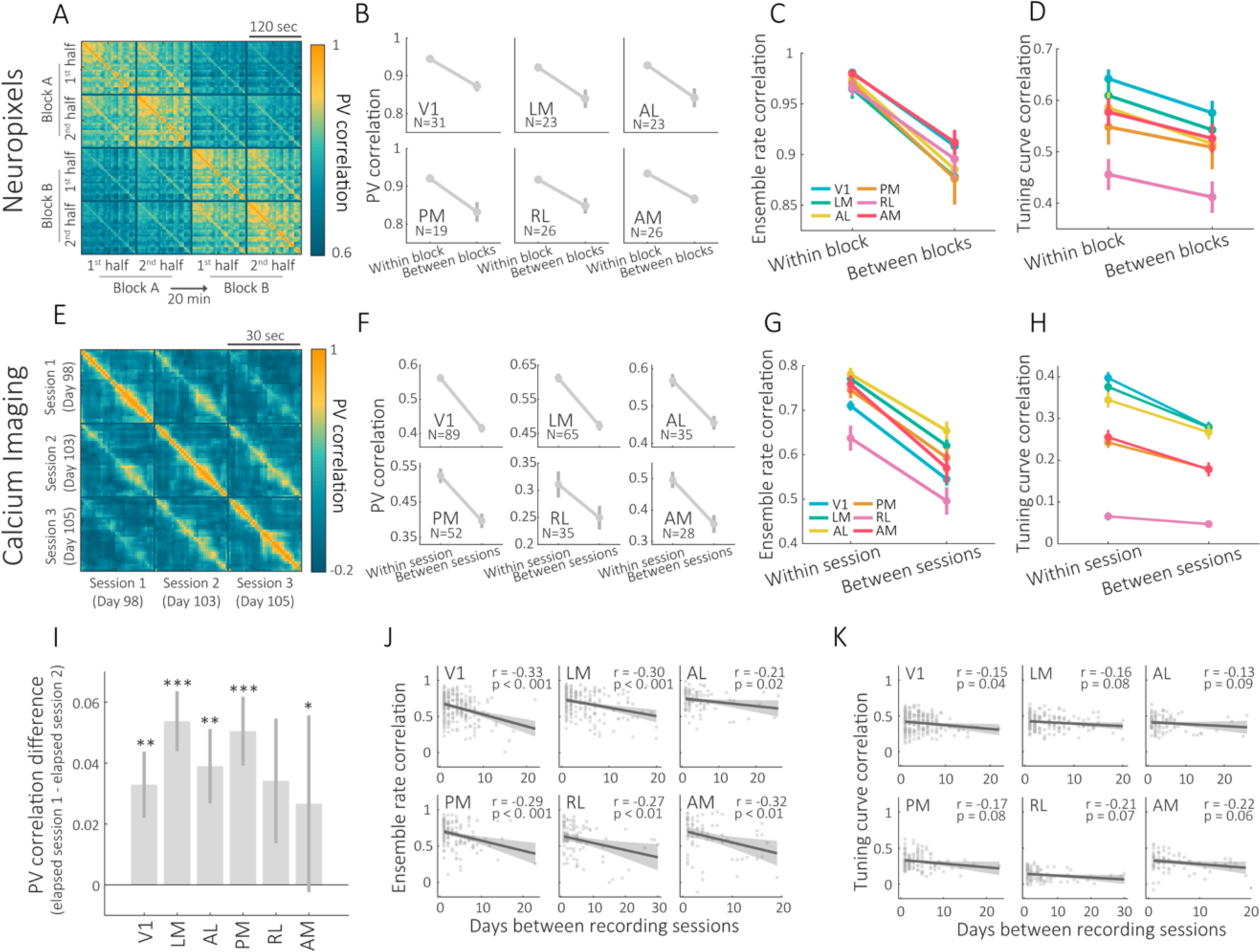
Visual representations gradually change over timescales of minutes-days. **A**. correlation between the 1^st^ (repeats 1-2) and 2^nd^ (repeats 3-5) halves of two different blocks of ‘Natural movie 3’ in a single visual area. The presented examples are the mean matrices across mice recorded in area LM with Neuropixels probes. **B**. PV correlation between the two halves of the same block (within block) and between halves of different blocks (between blocks) using the Neuropixels dataset (p < 10^−3^ for all areas, two-tailed Wilcoxon signed-rank test with Holm– Bonferroni correction). **C**. Ensemble rate correlation between the two halves of the same block (within block) and between halves of different blocks (between blocks) using the Neuropixels dataset (for all areas p < 10^−3^, two-tailed Wilcoxon signed-rank test with Holm–Bonferroni correction). **D**. Tuning curve correlation between the two halves of the same block (within block) and between halves of different blocks (between blocks) using the Neuropixels dataset (for all areas p ≤ 0.002, two-tailed Wilcoxon signed-rank test with Holm–Bonferroni correction). **E**. PV correlation between three different sessions from a single representative mouse recorded in V1 using two-photon Ca^2+^imaging. The correlation between temporally proximal sessions are higher relative to the correlation between two distal sessions. The age of the mouse (in days) is indicated in parenthesis. **F**. PV correlation between the two halves of the same session (within session) and between halves of different session (between sessions) (p < 10^−5^ for all areas, two-tailed Wilcoxon signed-rank test with Holm–Bonferroni correction). **G**. Ensemble rate correlation between the two halves of the same session (within session) and between halves of different sessions (between sessions) (p < 10^−5^ for all areas, two-tailed Wilcoxon signed-rank test with Holm–Bonferroni correction). **H**. Tuning curve between the two halves of the same session (within session) and between halves of different sessions (between sessions) (for all areas p < 10^−3^, two-tailed Wilcoxon signed-rank tests with Holm–Bonferroni correction). **I**. The difference between the PV correlation of temporally proximal sessions (the average correlation between sessions 1&2 and between sessions 2&3) and that of two distal sessions (the correlation between session 1&3) (V1 (Z = 3.35, p = 0.001), LM (Z = 4.64, p < 10^−4^), AL (Z = 2.85, p = 0.006), PM (Z = 3.92, p < 10^−3^), RL (Z = 1.38, p = 0.083), AM (Z = 1.99, p = 0.046), one-tailed Wilcoxon signed-rank test with Holm–Bonferroni correction; * p<0.05, ** p<0.01, ***p<0.001). **J**. Ensemble rate correlation as a function of the number of days between sessions. **K**. Tuning curve correlation as a function of the number of days between sessions. In panels F and G, each mouse is represented by 2-3 data points, corresponding to different unique intervals between sessions, with a regression line of ± CI of 95% (one-tailed Pearson’s correlation with Holm– Bonferroni correction). Data in panels B-E are mean ± SEM across mice.

**Figure 4.**
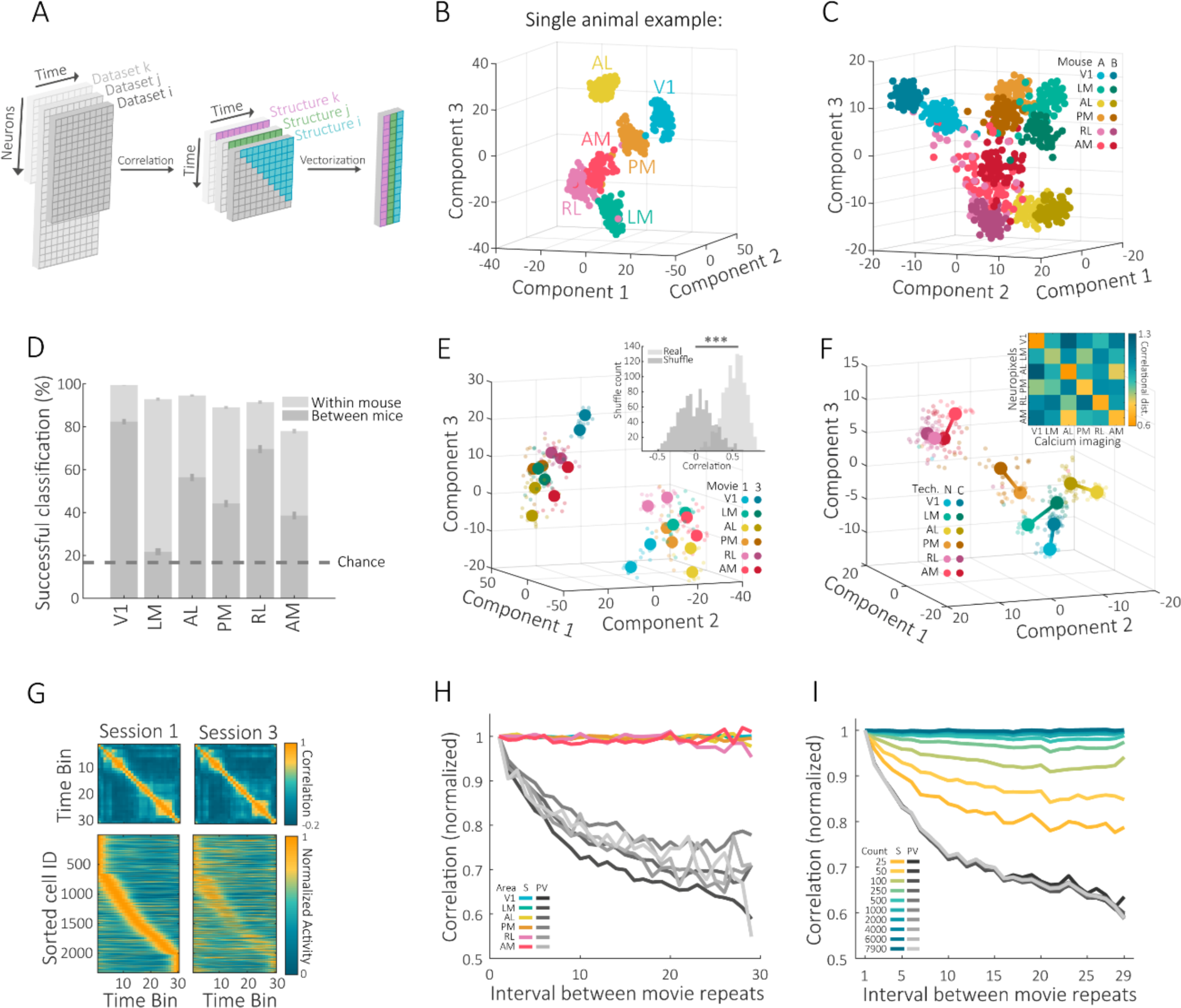
The internal structure of neuronal activity of each visual area is distinct, stereotypic, and stable over time. **A**. Workflow for the extraction of the internal structure from the population neuronal responses. Starting with a matrix (n x t) containing the mean neuronal activity in each temporal bin for each dataset (e.g., movie repeat, session, mouse, stimuli etc.). Correlating each temporal bin with the rest of the bins within a given dataset produces equally sized (t x t) matrices across datasets. Vectorising the upper half of these matrices produces vectors representing the internal structure (vector size = (t^2^-t)/2)). **B**. Dimensionality reduction (tSNE) applied to the internal structures of different visual areas from a single representative mouse recorded via Neuropixels. Each data point corresponds to an internal structure of a single ‘Natural movie 1’ repeat. **C**. Example for dimensionality reduction (tSNE) on the internal structures of ‘Natural movie 1’ produced from two Neuropixels ‘pseudo-mice’. Each data point corresponds to an internal structure of a single ‘Natural movie 1’ repeat. **D**. Percentage of successful classifications of the internal structures to their corresponding visual areas within and across Neuropixels pseudo-mice (mean ± SEM across n=100 pairs of pseudo-mice). **E**. Example for dimensionality reduction (tSNE) on the internal structures of ‘Natural movie 1’ and ‘Natural movie 3’ for two pseudo-mice. Each data point corresponds to an internal structure of a single movie repeat. Inset: For each pseudo-mouse and each natural movie, a correlation-distance matrix between the mean internal structure of each area was calculated. The correlations between the distance matrices of different movies across mice are significantly higher than those obtained in shuffled pseudo-mice (n=1000 pairs of pseudo-mice, p < 10^−20^, two-tailed Wilcoxon signed-rank test). **F**. Dimensionality reduction (tSNE) applied to the internal structures from different brain areas of two ‘pseudo-mice’ created using data from all the mice of each dataset. Each data point corresponds to an internal structure of a single repeat of ‘Natural movie 1’. Inset: a correlation-distance matrix between the median internal structure of each area across recording techniques. **G**. While the internal structure for ‘Natural movie 1’ in ‘pseudo-area AL’ is maintained across imaging sessions (top panels), the individual neurons whose activity patterns underlie the same internal structure drift across sessions (bottom panels). **H**. Correlation between the internal structures (colored lines) or the PVs (gray lines) as a function of the interval between repeats of ‘Natural movie 1’ from a single pseudo-mouse generated using data from all the mice in the Ca^2+^ imaging dataset. **I**. Correlation between the internal structures (colored lines) or the PVs (gray lines) as a function of the distance between repeats of ‘Natural movie 1’ in ‘pseudo-area V1’, plotted for varying numbers of neurons included in the analysis. Correlations in panels H and I were normalized to the value at the minimal interval between movie repeats.

To determine the degree to which visual representations change over timescales of tens of minutes, we analyzed the stability within and across blocks of movie presentations in both Neuropixels and Ca^2+^ imaging data. We found significantly higher correlations within a given block compared to between blocks in all measurements (PV correlation, ensemble rate correlation, and tuning curve correlation), brain areas and datasets, suggesting that visual representations change over the course of tens of minutes (Fig. 3A-D and Supp. Fig. 5A-D). Time-dependent changes in ensemble rate correlations were more pronounced than time-dependent changes in the cells’ tuning (Fig. 3C,D, Supp. Fig. 5C,D and Supp. Fig. 6A,B). Furthermore, the decline in ensemble rate correlations was continuous and proportional to the interval between blocks of different natural movies (Supp. Fig. 7A-F).

A major advantage of the Ca^2+^ imaging approach is the ability to record from the same neurons over multiple days, which allows examination of the long-term stability of neuronal representations (Sheintuch *et al*., 2017). The Ca^2+^ imaging dataset contains three imaging sessions per mouse, spanning multiple days (Fig. 3E). Previous time-lapse imaging studies in the hippocampus and posterior-parietal cortex have shown turnover in the ensemble of cells that are active on different imaging days (Ziv *et al*., 2013; Rubin *et al*., 2015; Driscoll *et al*., 2017). This turnover by itself can contribute to representational drift. Thus, we first took a conservative approach and included only cells that were active in both of the compared time points (either within a session or across sessions). Similarly to our observations within a given day, we found a significant representational drift across days in all measurements and brain areas (Fig. 3F-H). Notably, also across-days, changes in ensemble rate correlations were more pronounced than changes in the cells’ tuning (Fig. 3G,H and Supp. Fig. 6C). Furthermore, in most visual areas, the PV correlations between two proximal sessions were significantly higher than between two distal sessions, suggesting that representational drift is continuous over multiple sessions that occur on different days (Fig. 3I and Supp. Fig. 8A).

The distribution of the mean activity rates was similar across sessions in all visual areas, and there was also no difference in the PV correlation values between different pairs of subsequent sessions (i.e., the similarity between sessions 1 and 2 was comparable to the similarity between sessions 2 and 3), suggesting that representational drift over timescales of days is not a result of a gradual deterioration in neuronal activity or tuning (Supp. Fig. 8B,C).

To determine the effect of elapsed time, we compared the ensemble rate and tuning curve correlation between pairs of sessions separated by a different number of days. While ensemble rate correlations significantly decreased as a function of the number days between sessions in all visual areas, the tuning curve correlations showed only a modest trend (Fig. 3J,K). We also examined the extent to which representational drift is affected by the dynamic recruitment of cells to the representation on different days (Ziv *et al*., 2013; Rubin *et al*., 2015; Driscoll *et al*., 2017), by repeating our analyses with the pool of cells found active in at least one of the time points we compared (Supp. Fig. 8D). This analysis revealed an even more pronounced decline in the difference between subsequent sessions (Supp. Fig. 8E). Notably, time-dependent changes in visual representations were different across cortical layers in different areas (Supp. Fig. 9A,B), with layers 2/3 and 5 showing more stable representations than layers 4 and 6.

To what extent does the hierarchy of information flow across visual areas affect the stability of visual representations? To address this issue, we considered recent anatomical and functional studies that established a hierarchical structure of the mouse visual system (Harris *et al*., 2019; Siegle *et al*., 2019). In this hierarchy, the lateral geniculate nucleus (LGN) is at the bottom, and area AM is at the top. Therefore, in our analysis we compared the stability of thalamic areas (dorsal LGN and LP) and cortical areas (V1 and LM). Brain areas within these pairs are anatomically adjacent and show similar degree of tuning reliability to natural movies, but are distinct with respect to their level in the hierarchical structure of the visual system. We found that V1 was consistently less stable than the downstream area LM across all measured timescales (Supp. Fig. 10A-H), and likewise, LGN showed faster drift compared to the downstream area LP (Supp. Fig. 10A,B). Thus, our results do not support the hypothesis that lower visual areas are more stable over time than higher areas.

Our analyses so far have shown that the changes in visual representations occur across timescales that range from seconds to days. This raises the question of how could the visual system generate consistent perception despite representational drift and variability in neuronal responses. Recent studies in the hippocampus have shown that the structure of the relationship between neuronal population activity patterns remains stable over timescales of days, and is also stereotypic across mice (Rubin *et al*., 2019). Such a population-level representation may confer perceptual constancy in the face of changing coding properties of individual neurons. Thus, we next asked to determine the degree to which the structure of neuronal population activity is distinct for each visual area, stereotypic across individuals, and stable over time. To this end, we calculated for each area the PV for each time bin within a movie repeat and then calculated the correlations across all the PVs. This yielded a matrix (time by time) that represented the structure of similarities between representations (i.e. the ‘internal structure’ of neuronal population activity) of different time bins in the presented movie (Fig. 4A and Supp. Fig. 11A). We then applied dimensionality reduction to the vectors representing the internal structures for all movie repeats from all visual areas (see Methods). We found that the data was highly clustered, clearly separating between the different visual areas (Fig. 4B). This result suggests that the neuronal population activity of each visual area has a unique internal structure, and raises the possibility that the relationship among these structures is stereotypic.

It remains unclear, however, to what extent such an organization stems from the intrinsic functional properties of each brain area (Andermann *et al*., 2011; Glickfeld *et al*., 2013; Juavinett and Callaway, 2015; Murakami, Matsui and Ohki, 2017; Zatka-Haas *et al*., 2018), and to what extent it is susceptible to biases in the analysis (e.g., due to incidental differences in the coding properties of the sampled neurons or differential effects of the behavioral state on neuronal activity in different areas). To address these issues, we divided the dataset into two equal groups of mice, and then pooled together the data from each group to create two independent ‘pseudo-mice’, taking the same number of cells for each visual area in both pseudo-mice (Supp. Fig. 12A). Hence, the resultant pseudo-mice have an equal number of randomly-sampled neurons for all visual areas, with an order of magnitude more neurons per visual area compared to individual mice. Applying dimensionality reduction to the data from the two pseudo-mice revealed well-separated clusters (Fig. 4C and Supp. Fig. 11B) that correspond to the different visual areas, similarly to what we found in individual mice (Fig. 4B).

Next, we sought to quantify how distinct are the internal structures of different visual areas. To this end, we used a *k*-nearest neighbors algorithm to decode the identity of the recorded visual area based on the internal structure of single movie repeats, and found good classifications in all brain areas, both within and between pseudo-mice (Fig. 4D and Supp. Fig. 11C). The decoder’s performance within pseudo-mice was also higher than those of the same decoder in which we shuffled the identity of the neurons across visual areas (Supp. Fig. 11D-F), indicating that the performance of such decoder genuinely reflects the diversity in the coding properties across the different visual areas. Importantly, the internal structures of all visual areas (except of area LM) most closely resembled their equivalent structures across pseudo-mice (Fig. 4D and Supp. Fig. 11F,G), suggesting that the internal structure of each visual area is stereotypic. This was also true for the relationship between the internal structures of different brain areas across two different natural movies (Fig. 4E and Methods). Importantly, the relationship between the internal structures of the different areas was preserved across movies and pseudo-mice (Fig. 4E, inset and Supp. Fig. 12B). As another independent verification of these results, we constructed two pseudo-mice using datasets from the two recording techniques: one pseudo-mouse for the Ca^2+^ imaging data and another for the Neuropixels data (Fig. 4F). Consistent with our previous analyses, even across-recording techniques, all six visual areas were most similar to themselves across pseudo-mice (Fig. 4F, inset). Together, these analyses suggest that the different brain areas form a stereotypic relationship between their internal structures.

Finally, we examined whether the internal structure of neuronal population activity is more stable over time than the activity rates and tuning of individual neurons. Therefore, for each brain area we calculated the change in the correlations between the internal structures, and compared it to the change in the PV correlation as a function of the interval between movie repeats. While the PV correlations decayed with time, the correlations between the internal structures remained stable (Fig. 4G,H and Supp. Fig. 11H). The stability over time of the internal structure depended on the size of the neuronal population (Fig. 4I and Supp. Fig. 11I); including more cells in the analysis resulted in a more stable structure. Conversely, the change in PV correlation over time did not depend on the number of cells, consistent with a measurement that treats cells independently. Overall, our results suggest that the internal structure of neuronal population activity of each visual area is distinct, stereotypic, and stable across time.

## Discussion

Here, we used large-scale electrophysiological and optical imaging data to systematically study the stability of neuronal encoding of complex visual stimuli across different areas of the visual cortex (Siegle *et al*., 2019; de Vries *et al*., 2020). Our results demonstrate the existence of a representational drift over timescales of minutes to days in all studied brain areas. Similarly to previous findings in the hippocampus, we found that drift in visual representations is primarily driven by changes in cells’ activity rates, while their tuning changes to a lesser degree (Ziv *et al*., 2013; Rubin *et al*., 2015). Surprisingly, our analysis does not support the hypothesis that primary (or lower) sensory areas should display more stable coding than downstream (higher) areas (Haak, Morland and Engel, 2015; Haak and Mesik, 2016). If anything, our analysis shows that the coding stability of some cortical (V1 and V2) and subcortical (LGN and LP) areas exhibit an opposite trend with respect to their hierarchy. These results are in line with a recent study in the human visual cortex that demonstrated, using functional MRI, a non-monotonic relationship between plasticity and hierarchical level (Haak and Beckmann, 2019). We further show that the structure of the relationship between neuronal population activity patterns is stereotypic across mice and stable over time, pointing to a possible network mechanism that can reliably preserve visual information despite drift in the coding properties of individual neurons.

Our work joins a number of longitudinal studies that quantified coding stability in the visual cortex and adds to these studies in several important aspects (Montijn *et al*., 2016; Rose *et al*., 2016; Ranson, 2017; Jeon *et al*., 2018). While previous work focused on V1, our analysis encompasses six different visual areas (along with two sub-cortical structures), allowing a direct comparison between them. Whereas earlier studies used mostly simple synthetic stimuli (e.g., static or moving gratings), our study characterizes neuronal responses to natural movies, a complex and more ethologically relevant visual stimulus. While most previous investigations focused on L2/3 neurons, in this work we were able to analyze recordings from different cortical layers. Previous studies used either electrophysiological recordings or two-photon Ca^2+^ imaging and had different experimental schedules, which precluded a direct comparison of findings made using the different techniques. Here, neuronal responses to the exact same stimuli were recorded using both electrophysiology and Ca^2+^ imaging, which allowed us to validate the results and control for biases specific for each technique (see below). The datasets from both recording techniques also allowed us to study coding stability across different timescales (seconds to days).

In this work we deconstructed a commonly applied measure of coding stability (PV correlations)(Leutgeb *et al*., 2005) into two measures: ensemble rate correlation and tuning curve correlation. By doing so, we found that similarly to findings in hippocampus (Ziv *et al*., 2013; Rubin *et al*., 2015), time-dependent changes in activity rates are more dominant than changes in the cells’ tuning curve. These two measures presumably capture changes in different aspects of neuronal physiology, such as excitability (rate) and synaptic connectivity (tuning). Our findings that tuning curve correlations change over time to a lesser extent than do ensemble rate correlations are in line with previous work that found tuning to visual stimuli to be relatively stable (Montijn *et al*., 2016). However, both measures revealed significant changes over time across most brain areas, which highlights the fact that coding stability is a multi-dimensional property. Whether time-dependent changes along such different dimensions of coding stability have a qualitatively different impact on the function of the visual system remains to be studied.

Recent studies suggest that, in some brain areas, representational drift could have a beneficial role. For example, in the hippocampus and lateral entorhinal cortex, representational drift may support the encoding of time in episodic memory (Manns, Howard and Eichenbaum, 2007; Ziv *et al*., 2013; Cai *et al*., 2016; Tsao *et al*., 2018). By encoding each experienced event in a unique manner, representational drift can help deal with catastrophic interferences, or serve as a means to continuously update previously learned associations (Wiskott, Rasch and Kempermann, 2006; Eichenbaum, 2017). Overall, although it is currently unclear whether the functionality of the sensory cortex could benefit from representational drift, we cannot rule out such an option.

In our analysis, we were able to demonstrate the existence of representational drift, whereas previous work emphasized variability (Montijn *et al*., 2016; Ranson, 2017; Jeon *et al*., 2018). Notably, in some of these studies, drift could not be determined because only two time points were compared. However, irrespective of whether changes in the coding of visual information are due to drift or variability, the visual system must cope with them to generate consistent perception. It has been suggested that a system that carries a high-dimensional distributed code may maintain its functionality under representational drift by either confining the drift to the null space of the code, or via a compensatory plasticity of the downstream reader (Rule, O’Leary and Harvey, 2019; Rule *et al*., 2020). In both cases the similarities across representations of different stimuli are expected to be somewhat conserved over time, even under a significant change in the representations themselves. Here, we took advantage of the large number of mice in Allen Brain Observatory datasets, and generated pseudo-mice, which consisted of between ten-to-hundred-fold more neurons per brain area than in each real mouse. This approach allowed us to portray and analyze the internal structure of neuronal population activity, while avoiding biases due to the size of the sampled population of neurons, noise, or variability in the behavioral state across mice. Here, again, obtaining the same results using data from the two recording techniques provided additional validation of this analysis approach. We found that different visual areas had distinct internal structures and that the internal structure for a given brain area was similar across mice. We also demonstrated that the internal structure was stable over time across all visual areas, and that its stability depends on the number of cells included in the analysis. These results are consistent with several recent studies that showed that a stable structure (manifold) of population activity underlies a stable behavior (Rubin *et al*., 2019; Bolding *et al*., 2020; Gallego *et al*., 2020; Pashkovski *et al*., 2020). Our results are also consistent with a recent study in V1 that showed that high-dimensional population codes are more stable than low-dimensional population codes (Montijn *et al*., 2016). In line with that work, our population-level analysis hints at the possibility that the sheer size of the neuronal population in a given area contributes to the robustness of the visual representation to drifting responsiveness of individual neurons.

Measuring coding stability is challenging because various factors could affect longitudinal recordings in a way that could lead to the appearance of drift, even if the neuronal activity itself is stable. In electrophysiological recordings, movement of the electrode inside the tissue can cause some units to disappear and others to appear, which will show in the analysis as variable or drifting responses. Biological reactions to the implanted probes, such as scaring and inflammation around the electrode, could likewise lead to gradual deterioration of the recorded signal. Given these issues, traditional electrophysiological recording techniques cannot be reliably used for longitudinal studies over timescale longer than ∼1-2 days. In this work, we analyzed electrophysiological data recorded in head-fixed mice using Neuropixels probes (Jun *et al*., 2017), and focused on changes in neuronal responses over timescales of tens of minutes (Fig. 2 and Fig. 3A-D), during which recording instability is less likely to play a role. Moreover, several control analyses suggest that our results are not due to recording instability: (1) We found similar effects within a block even when we included in our analysis only units that were highly stable across blocks (Supp. Fig. 4). (2) We found continuous changes in the ensemble activity across blocks (over > 1 hour), even when we included in the analysis only units that showed high tuning curve correlation across blocks and across two different movies (Supp. Fig. 7). (3) We found similar results in data recorded using two-photon Ca^2+^ imaging (Supp. Fig. 1D-F and Supp. Fig. 5A-D).

Notably, while two-photon Ca^2+^ imaging is suitable for time-lapse recordings from populations of the same neurons over weeks and even months, variability in the output of cell detection or cell registration algorithms may lead to upward bias in the turnover-rates in the identity of cells that are active in different sessions (Sheintuch *et al*., 2017). To address this issue, we focused our analysis on cells that were active in each pair of time points compared. Furthermore, we found no evidence of a gradual time-dependent deterioration of the neural code (Supp. Fig. 8B,C). Overall, although we cannot completely rule out any contribution of recording instability to our results, we find this possibility to be unlikely.

Gradual alterations in the code could also result from changes in animal behavior or attention (Niell and Stryker, 2010; Keller, Bonhoeffer and Hübener, 2012; Ayaz *et al*., 2013; Polack, Friedman and Golshani, 2013; Erisken *et al*., 2014; Ruff and Cohen, 2014; Vinck *et al*., 2015; Mineault *et al*., 2016; Dipoppa *et al*., 2018; Musall *et al*., 2019). Indeed, mouse running speed and pupil area declined during the first five (out of 30) movie repeats, possibly reflecting changing attention or arousal within a block of movie repeats. However, our findings show that drift rates remain similar also when the first eight movie repeats were excluded from the analysis (Supp. Fig. 2). Thus, representational drift occurs even in the absence of an overt sign of changes in the behavioral or cognitive state.

In summary, the availability of large-scale recording datasets by the Allen Brain Observatory make it possible to conduct an in-depth investigation of coding stability across different visual cortical areas and a wide range of temporal scales. Here, we conducted such an investigation and found the existence of representational drift in all studied areas. The available two-photon imaging data allowed a longitudinal investigation over timescales of days, but since different mice had different intervals between the three recording sessions, we could not quantify the lifetime of visual representations in different cortical areas. Future studies that consist of multiple equally spaced sessions over many days and weeks are needed to fully address this issue. Currently, the degree to which our findings could be generalized to other sensory cortical areas is unknown; still, our findings across multiple cortical areas imply that representational drift is an inherent property of neural networks, and that population-level organization of information could contribute to robust, time-invariant representations despite drifting or variable coding at the level of individual neurons.

## Methods

### Data curation

We analyzed data from the publicly available Allen Brain Observatory: two-photon calcium imaging(de Vries *et al*., 2020) and electrophysiology (Neuropixels) datasets (Siegle *et al*., 2019; de Vries *et al*., 2020). We used the default functions in AllenSDK package to download the raw Neurodata Without Borders (NWB) files containing the neuronal and behavioral data from the relevant experiments. Their full data collection methodology can be found in the white paper (https://observatory.brain-map.org/visualcoding). In the calcium imaging dataset, 216 transgenic mice expressing GCaMP6f in laminar-specific subsets of cortical pyramidal neurons underwent intrinsic signal imaging to map their visual cortical regions before cranial windows were implanted above the desired visual region. Mice were habituated to head fixation before the three imaging sessions, in which they were shown a battery of natural scenes, natural movies, locally sparse noise, or gratings. In the Neuropixels dataset, 30 C57BL/6J wild-type mice and 28 mice from three transgenic lines (N = 8 Pvalb-IRES-Cre x Ai32, N = 12 Sst-IRES-Cre x Ai32, and N = 8 Vip-IRES-Cre x Ai32) were implanted with up to six Neuropixels silicone probes each. The dataset contains simultaneous recordings from up to 8 cortico-thalamic visual areas (as well as nearby regions, such as hippocampus). During each recording session, mice passively viewed a battery of natural and artificial stimuli, depending on their experimental group.

### Data analysis

Analysis was carried out using both AllenSDK package default functions (for data curation) and custom-written MATLAB scripts (for data analysis). In the Ca^2+^ imaging dataset, we analyzed all available excitatory Cre-lines, including all layers and brain areas. The dataset is structured into ‘experiment containers’ that group recordings from three different imaging sessions of the same field of view. We considered each such container as an individual mouse. We included only mice that passed a fixed criterion of at least 20 recorded cells in the compared time points. Specifically, in the within-block and between-days analysis (Supp. Figs. 1-2 and Fig. 3, respectively), we included only mice that had at least 20 recorded cells in each of the three imaging sessions. In the between-blocks analysis (Supp. Fig. 5A-D), we included only mice that had at least 20 recorded cells within the same session (‘Session A’). In the Neuropixels dataset, we used the AllenSDK package default functions to retrieve the relevant unit’s identity according to their corresponding manually labeled brain areas. These units passed a set of quality criteria to ensure their validity. We then included in all analyses (except those presented in Fig. 4) only data from areas with at least 15 recorded units in each mouse.

### Detection of Ca^2+^ events

Neuropil-corrected fluorescence change (ΔF_(t)_/F_0_) traces for each cell were extracted using automated, structural region of interest (ROI) based methods (see Allen Institute white paper for details, https://observatory.brain-map.org/visualcoding). We performed no further preprocessing on ΔF_(t)_/F_0_ traces after downloading them through the AllenSDK. We identified Ca^2+^ events by searching each trace for local maxima that had a peak amplitude higher than four times the entire trace absolute median while including only the frames that showed an increase in Ca^2+^ transients relative to their previous frame. All the ΔF_(t)_/F_0_values in the frames that passed the assigned filters were set to the value of 1 and the rest were set to a value of 0.

### Registration of cells across sessions

We used each cell’s match labels across sessions as was calculated and provided by Allen Brain Institute in each experiment’s NWB file. Briefly, an algorithm that combines the degree of spatial overlapping and closeness between the ROIs of different cells was used to create a unified label for each cell across all three sessions. The full registration procedure appears in de Vries et al. (2020).

### Visual stimuli

In our analysis we only used ‘Natural movie 1’ (30-second clip) and ‘Natural movie 3’ (120-second clip) stimuli from the Allen Brain Observatory paradigm. In the calcium imaging dataset, ‘Natural movie 1’ was presented across all three imaging sessions (ten repeats per session). ‘Natural movie 3’ was presented only in one of the sessions (session A), with ten repeats spanning two blocks (five repeats in each block). In the Neuropixels dataset, ‘Natural movie 1’ was presented with either 60 repeats spanning two blocks (30 repeats in each block) for the ‘Functional connectivity’ group, or with 20 repeats spanning two blocks (10 repeats in each block) for the ‘Brain observatory 1.1’ group. ‘Natural movie 3’ was presented with ten repeats spanning two blocks (five repeats in each block) only for the ‘Brain observatory 1.1’ group.

### Population vector correlation

To determine the level of similarity between visual representations of the same stimulus on different presentations, we calculated for each mouse the population vector correlation between pairs of different movie repeats. First, we divided each movie repeat into 30 equal time bins (each bin spanning 1 sec in ‘Natural movie 1’ and 4 secs in ‘Natural movie 3’). Then, for each temporal bin, we defined the population vector as the activity rate for each cell/unit. We calculated the Pearson’s correlation between the population vector (PV correlation) in one repeat with that of all temporal bins in another movie repeat, and averaged the correlations over all pairs of corresponding time bins. For the between-blocks analysis, we created two mean PVs for each of the two blocks; one PV from the first two ‘Natural movie 3’ repeats (repeats 1-2), and a second PV from the last three repeats (repeats 3-5). We than calculated the Pearson’s correlation across all four vectors of both blocks and measured the difference between PV correlations within blocks and across blocks. The mean correlations between the two PVs of the same blocks capture the ‘within-block’ stability, and the mean correlations between different blocks, capture the ‘between-blocks’ stability. The between-days analysis is similar to that of between blocks with minor changes: For each ‘Natural movie 1’ session, two mean PVs were calculated, one vector from the first five ‘Natural movie 1’ repeats (repeats 1-5) and a second vector from the last five ‘Natural movie 1’ repeats (repeats 6-10). We then calculated the Pearson’s correlation between each pair of PVs, including in the PVs of only the cells that were active in both time points, and measured the difference in PV correlations within blocks and across blocks. The mean correlations between the two PVs of the same session capture the ‘within-session’ stability, and that of different blocks, the ‘between-sessions’ stability. For the analysis shown in Fig 3 I and Supp. Fig 8C, PV correlations were calculated after averaging the activity rate of each individual neuron over all movie repeats in a given session.

### Ensemble rate correlation

To quantify the similarities in activity patterns between different presentations of the same stimulus (regardless of the specific tuning of each neuron), we calculated for each mouse the ensemble rate correlation between pairs of different movie repeats. First, we calculated the overall activity rate for each neuron in each movie repeat. We then calculated for each pairs of movie repeats the ensemble rate correlation as the Pearson’s correlation between their vectors of activity rates. As in the PV correlation analysis, the differences in ensemble rate correlation for within and between blocks (or sessions) were calculated after averaging the activity rates of individual neurons over the first and second halves of movie repeats in each block (or session). For the analysis shown in Fig 3J and Supp. Fig 8A, ensemble rate correlations were calculated after averaging the activity rate of each individual neuron over all movie repeats in a given session.

### Tuning curve correlation

To quantify the similarities in the tuning preference of individual neurons across different presentations of the same stimulus (regardless of changes in activity rates), we calculated for each neuron the tuning curve correlation between different movie repeats. As in the PV correlation analysis, we first divided each movie repeat into 30 equal time bins (each bin spanning 1 sec in ‘Natural movie 1’ and 4 secs in ‘Natural movie 3’). Then, for each neuron, we defined the tuning curve as the mean activity rate in each temporal bin within the movie. We calculated the Pearson’s correlation between the tuning curve of each individual neuron in one movie repeat and that of the same neuron in another movie repeat, and used the median value across all neurons to capture the central tendency of the entire population. As in the PV correlation analysis, the differences in tuning curve correlation for within and between blocks (or sessions) were calculated after averaging the activity rates of individual neurons for the first and second halves of movie repeats in each block (or session). For the analysis shown in Fig 3K and Supp. Fig 8A, tuning curve correlations were calculated after averaging the activity rate of each individual neuron over all movie repeats in a given session. Due to the sparseness in neuronal responses in the Ca^2+^ imaging dataset, we used the mean value across all cells (instead of the median) when computing the tuning curve correlation between individual movie repeats (Supp. Fig. 1F and Supp. Fig. 10D).

### Time-lapse decoding analysis

(related to Supp. Fig. 3). We used a *k*-nearest neighbors classifier with K=1 to decode the time bin at a given movie repeat based on the population vectors of a preceding repeat using the Euclidean distance between the response vectors. The performance of the decoder was defined as the percentage of correct classifications out of the 30 time bins for each pair of movie repeats.

### Internal structure of neuronal population activity

Similar to the PV correlation analysis, we divided each movie repeat into 30 equal time bins and calculated the population vector for each time bin, yielding a matrix of 30 by the number of recorded neurons. Then, we calculated the Pearson’s correlation across all vectors of the same movie repeat, resulting in a symmetric 30-by-30 matrix. This matrix represented the structure of similarities across the population activity patterns at all different time bins of the presented movie. We defined this matrix of similarities as the ‘internal structure of neuronal population activity’ (or ‘internal structure’). Since this structure no longer holds the identities of individual neurons, it is possible to measure the resemblance between structures extracted from different datasets (e.g., movie repeats, natural movies, areas, mice, etc.) without relying on the ability to record from the same cells or requiring equal numbers of cells across measurements.

### Pseudo-mice and shuffled pseudo-mice

(related to Fig. 4 and Supp. Fig. 11). To reduce the effect of incidental differences in the coding properties of the sampled neurons on our ability to capture the true internal structure of each of the studied areas, we constructed ‘pseudo-mice’, which are a pooling of cells recorded from different mice of the same dataset (Supp. Fig. 12A). To create two independent pseudo-mice (i.e. pseudo-mice that have no overlap in their source of neuronal activity), we first randomly split the complete Neuropixels dataset of 58 mice into two non-overlapping groups of 29 mice. Each mouse in each group contained the neuronal activity recorded from 1-6 brain areas. Pooling all the cells/units from each brain area across all mice of the same group yielded six distinct sets of neurons (one per area) for each of the two pseudo-mice (12 pseudo-areas in total). Since there is variability in the number of recorded areas and cells across mice, the pooling procedure resulted in different number of cells in each of the pseudo-areas. To ensure that differences between the internal structures of different areas did not stem from the size of the recorded neuronal population, we randomly subsampled an equal number of cells from the entire population of each area. The exact number of subsampled cells was determined based on the pseudo-area with the lowest number of cells among both pseudo-mice. To verify the uniqueness of the internal structure of each area, the analysis was compared to complementary ‘shuffled pseudo-mice’ that were created by the random redistribution of all the cells across areas in each of the pseudo-mice.

### Within and across ‘pseudo-mice’ decoding

(related to Fig. 4 and Supp. Fig. 11). Since the two groups of mice in the Neuropixels dataset (‘Brain Observatory 1.1’ and ‘Functional connectivity’) were presented with different number of ‘Natural movie 1’ repeats (20 and 60, respectively), the analysis was performed using only the first 20 repeats. For the within-pseudo-mouse decoding, we used a *k*-nearest neighbors classifier with K=1 to decode the area of origin for a single internal structure (representing a single movie repeat) based on the cosine distances to rest of the internal structures of the same pseudo-mouse. To evaluate the performance of the decoder, we calculated the percentage of internal structures that were correctly classified to their corresponding area within each of the two pseudo-mice and later averaged across them. For the between-pseudo-mice decoding, we used a *k*-nearest neighbors classifier with K=1 to decode the area of origin for a single internal structure of a given pseudo-mouse based on the cosine distances to the internal structures of the other pseudo-mouse. To evaluate the performance of the decoder, we calculated the percentage of internal structures that were correctly classified to their corresponding area across pseudo-mice. The analysis was repeated 100 times to obtain representative results across different realizations of pseudo-mice (different realizations of dividing the mice population into two random subsets) and was compared to the results obtained when using shuffled pseudo-mice.

### Relationship between internal structures of different brain areas across natural movies

(related to Fig. 4E and Supp. Fig. 12B). Since the two movies are of different lengths (30 seconds and 120 seconds) and were presented different number of times (20 and 10 repeats), the analysis was performed on the first 30 seconds and first 10 movie repeats of both ‘Natural movie 1’ and ‘Natural movie 3’ using only the mice from the Neuropixels ‘Brain observatory 1.1’ group (the group that was presented with both movies). To measure the degree to which the relationship between the internal structures of different areas is conserved across different movies and pseudo-mice, we first calculated for each brain area of each pseudo-mouse the internal structure per movie repeat. We then averaged the internal structures over all movie repeats to create 24 mean internal structures (6 areas x 2 natural movies x 2 pseudo-mice). Then, for each natural movie within a given pseudo-mouse, we calculated the Pearson’s correlation matrix across the internal structures of all areas. This procedure yielded four matrices (2 natural movies x 2 pseudo-mice), each symmetric and 6-by-6 in size (across all areas). Finally, we calculated the Pearson’s correlation between the matrices (using the vectorization of the upper half of each matrix) of different natural movies and different pseudo-mice, and averaged across the two comparisons (correlation between ‘Natural movie 1’ in pseudo-mouse A with that of ‘Natural movie 3’ in pseudo-mouse B, and vice versa). The analysis was repeated 1000 times to obtain representative results across different realizations of pseudo-mice (different realizations of dividing the mice population into two random subsets), and was compared to the results obtained when using shuffled pseudo-mice.

### Similarity between internal structures of the same brain areas across recording technologies

(related to Fig. 4F). In this analysis, we used ‘Natural movie 1’ data from all mice of the Neuropixels dataset and all mice of the two-photon Ca^+2^ imaging dataset to create two pseudo-mice, one for each of the recording techniques. Since different mice in the Neuropixels dataset were presented with different number of movie repeats, 20 repeats in the ‘Brain Observatory 1.1’ group and 60 repeats in the ‘Functional connectivity’ group, we used only the first 20 repeats for ‘Functional connectivity’ group. First, we calculated for each brain area of each pseudo-mouse the internal structure per movie repeat. We then calculated the median internal structures over all movie repeats to create 12 representative internal structures (6 areas x 2 pseudo-mice). Finally, we normalized (z-score) the internal structures within each pseudo-mouse and calculated the Pearson’s correlation distance matrix across areas of the two pseudo-mice (Neuropixels pseudo-mouse and Ca^2+^ pseudo-mouse).

### Internal structure stability

(related to Fig. 4H). First, we created a single pseudo-mouse from all the ‘Natural movie 1’ data of the Ca^+2^ imaging dataset using only the cells that were active in all three recording sessions. Then, for each area, we calculated both the population vectors for all time bins and internal structure across all time bins for each of the 30 movie repeats. Lastly, we calculated the Pearson’s correlations for both measurements across all pairs of movie repeats and calculated the change in correlations as a function of the interval between movie repeats. Both measurements were normalized to the value of the smallest interval between movie repeats.

### Temporally shuffled internal structure

(related to Supp. Fig. 11H). To verify that the internal structure used in Fig. 4H contain information beyond the similarities between subsequent time bins, we calculated the internal structure stability, defined as the mean Pearson’s correlation between all pairs of internal structures across all 30 movie repeats, and compared it to the values obtained after performing a random cyclic temporal shuffling of the same internal structures. The analysis was repeated 100 times to obtain representative results across different shuffles.

### t-distributed Stochastic Neighbor Embedding

(t-SNE; related to Fig. 4 and Supp. Fig. 11). For visualizing the relationships between internal structures, the vectors of pairwise correlations across activity patterns were embedded in three dimensions using t-SNE (Maaten and Hinton, 2008; Maaten, 2014). We used the exact tSNE algorithm with similar embedding settings for all visualizations (cosine distance metric, using 10 PCA components, exaggeration 4 (default), and learning rate 500 (default)). The perplexity (effective number of local neighbors of each point) was chosen for each visualization manually based on multiplication of the minimal number of movie repeats used in the analysis (30, 60, 20 and 20 for Fig. 4 B,C,E and F, respectively). Note that embedding in the reduced space is used only for visualization of the relationships between internal structures, the analysis shown in Fig 4 D-F and Supp. Fig. 11 were performed on the pairwise distances between the original (non-reduced) vectors of internal structures as described in other sections of the Methods.

### Statistical analysis

All statistical details, including the specific statistical tests, are specified in the corresponding figure legends. In general, non-parametric Wilcoxon rank sum tests (unpaired data), signed rank tests (paired data), or parametric two-way mixed model ANOVAs (with Greenhouse-Geisser correction for sphericity assumption violation) were performed and corrected for multiple comparisons (using Holm–Bonferroni method), unless otherwise noted. A one-sided Pearson’s correlation coefficient was used to estimate the effect of elapsed time on ensemble rate and tuning curve stability. In all tests, significance was defined at α = 0.05. Aside from mice with low number of recorded cells (see ‘Data analysis’ section in the Methods), no neural data was excluded from analysis. All statistical analyses were conducted using MATLAB 2017b (Mathworks).

## Acknowledgments

Y.Z. is a CIFAR-Azrieli Global Scholar in the Brain, Mind & Consciousness program, and the incumbent of the Daniel E. Koshland Sr. Career Development Chair. Y.Z. is supported by grants from the Abraham and Sonia Rochlin Foundation, the Hymen T. Milgrom Trust, the Israel Science Foundation (grant 2113/19), the Human Frontier Science Program, and the European Research Council (ERC-StG 638644). We thank Timothy O’Leary, Ofer Yizhar, Rafi Malach, Ivo Spiegel, Michal Rivlin, Jerome Lecoq, Michael Rule, Meytar Zemer, Liron Sheintuch, Eyal Bitton, Maya Salomon, Alice Eldar, Ofer Givton, and Noa Eren for helpful advice and comments on the manuscript.

---

**Supplementary Figure 1.**
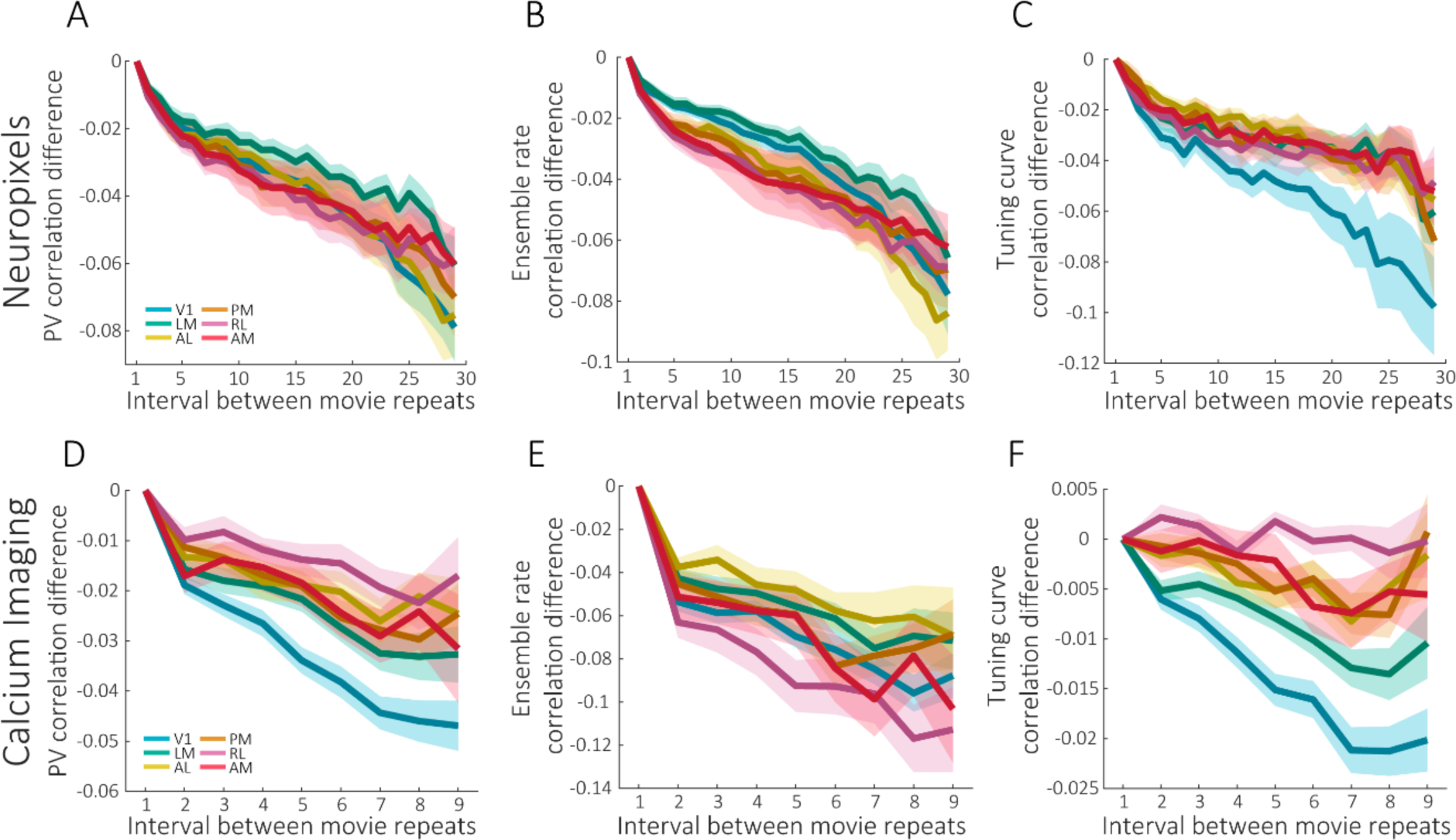
Representational drift over timescales of seconds-minutes: comparison between Neuropixels and Ca^2+^ imaging datasets. **A, D**. Difference in PV correlation as a function of the interval between movie repeats for six visual cortical areas. **B, E**. Difference in the ensemble rate correlation as a function of the interval between movie repeats for six visual cortical areas. **C, F**. Difference in the tuning curve correlation as a function of the interval between movie repeats for six visual cortical areas. Data in panels A-C are mean ± SEM across mice from the Neuropixels ‘Functional connectivity’ group. Data in panels D-F are mean ± SEM across mice from the Ca^2+^ imaging dataset. Correlations were scaled by subtracting the correlation value at the minimal interval between movie repeats.

**Supplementary Figure 2.**
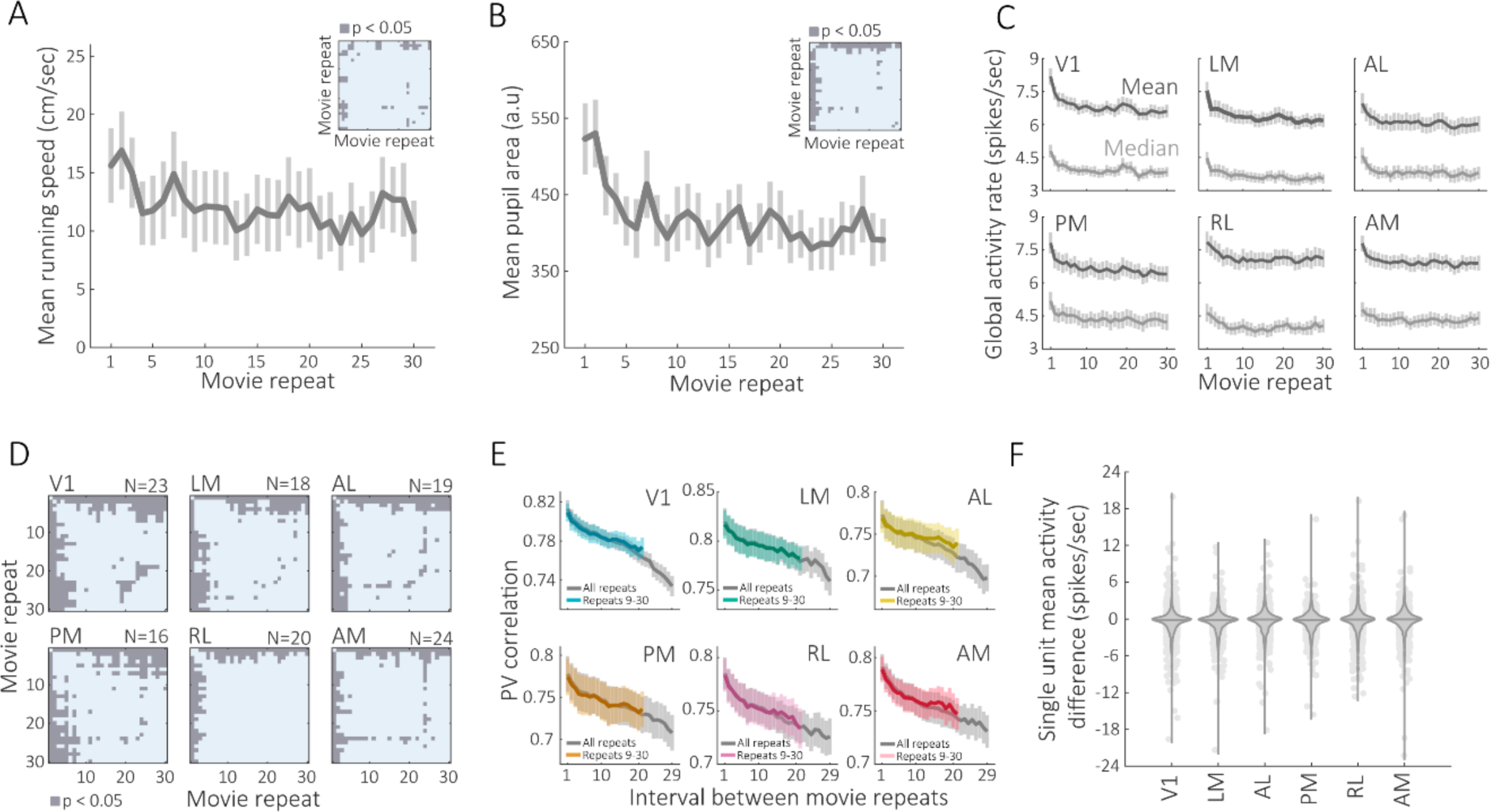
Representational drift is not driven by changes in behavioral state and global activity rates. **A**. Mean running speed for each movie repeat across animals. **B**. Mean pupil area for each movie repeat across animals. Insets in A and B indicate a significant difference between the repeats (paired t-test, p<0.05, two-tailed, without correction for multiple comparisons). **C**. Mean and median activity rates for each movie repeat across animals and brain areas. **D**. Testing the differences in mean activity rates between pairs of movie repeats for the data presented in panel C. Gray entries indicate a significant difference in activity rates between the repeats (paired t-test, p<0.05, two-tailed, without correction for multiple comparisons). **E**. Mean population vector correlation as a function of the interval between movie repeats, performed on a subset of movie repeats (repeats 9 −30; colored lines). PV correlations of this subset of the data gradually declined with the interval between movie repeats, similarly to the PV correlations of the full dataset (gray lines) from all movie repeats; The difference in PV correlations between the interval of one movie repeat and that of 21 movie repeats was significant for all areas (p ≤ 0.007, two-tailed Wilcoxon signed-rank test with Holm–Bonferroni correction). **F**. Distribution of mean activity difference of single units across areas. For each unit, the mean activity rates (spikes/sec) of repeats 2-6 was subtracted from the mean activity of repeats 26-30. The first repeat was removed from the analysis since it is characterized with non-representative activity patterns. Data in all panels are mean± SEM across mice from the Neuropixels ‘Functional connectivity’ group.

**Supplementary Figure 3.**
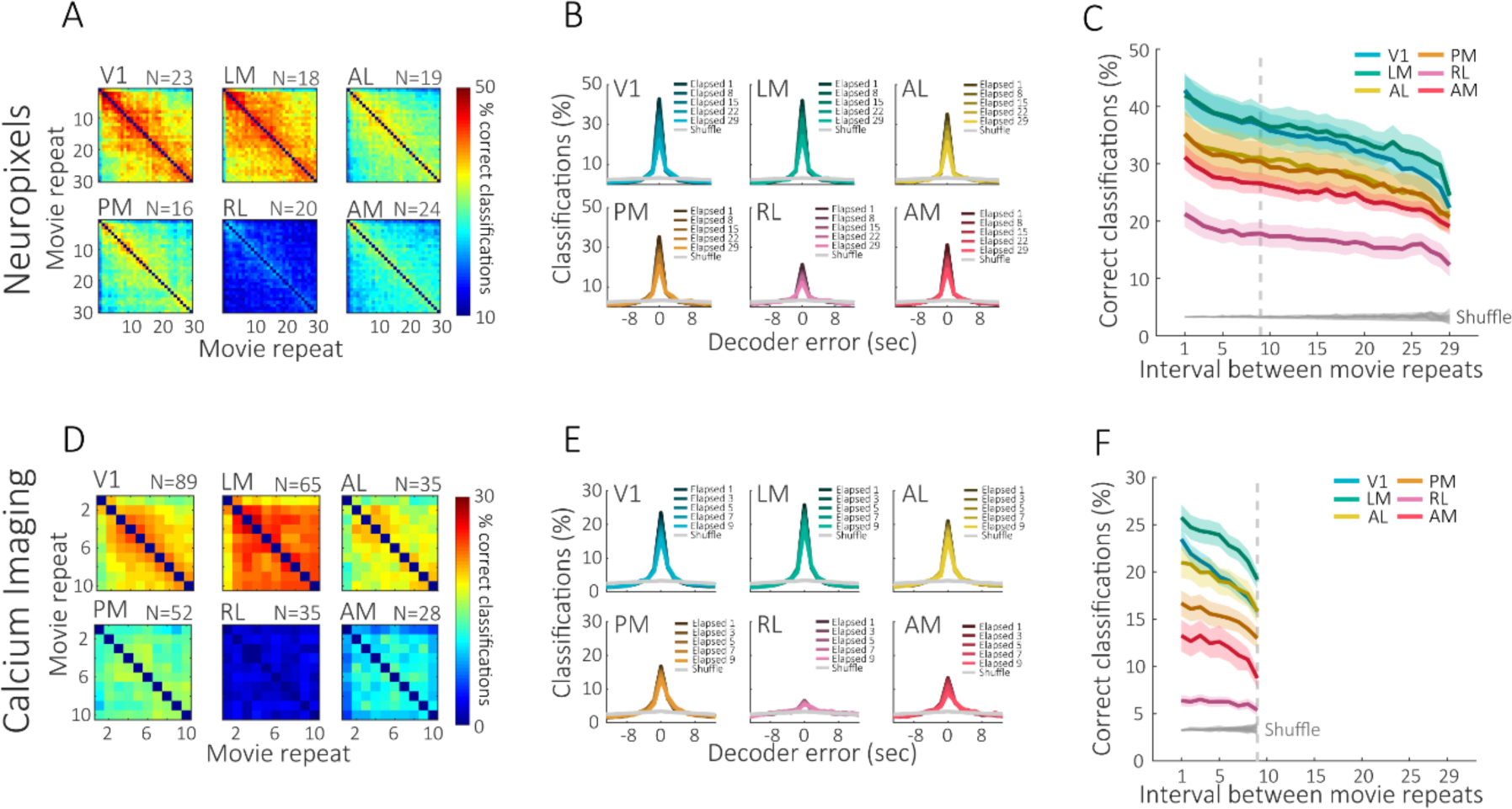
Time-lapse decoding analysis reveals representational drift across visual areas in both Neuropixels and Ca^2+^ imaging datasets. **A, D**. Mean percentage of correct classifications of the decoder (*k*-nearest neighbors) between repeats of ‘Natural movie 1’. **B, E**. Mean percentage of decoder classifications between movie repeats as a function of decoder error. Dark colors represent a short interval and light colors represent a long interval between the train and test data. **C, F**. Percentage of correct classifications as a function of the interval between the train and test movie repeats; the difference in correct classifications between the minimal and maximal interval of movie repeats was significant for all areas (p ≤ 0.011, two-tailed Wilcoxon signed-rank test with Holm–Bonferroni correction). Panels A-C, Neuropixels dataset. Panels D-F, Ca^2+^ imaging dataset. Data are mean± SEM across mice.

**Supplementary Figure 4.**
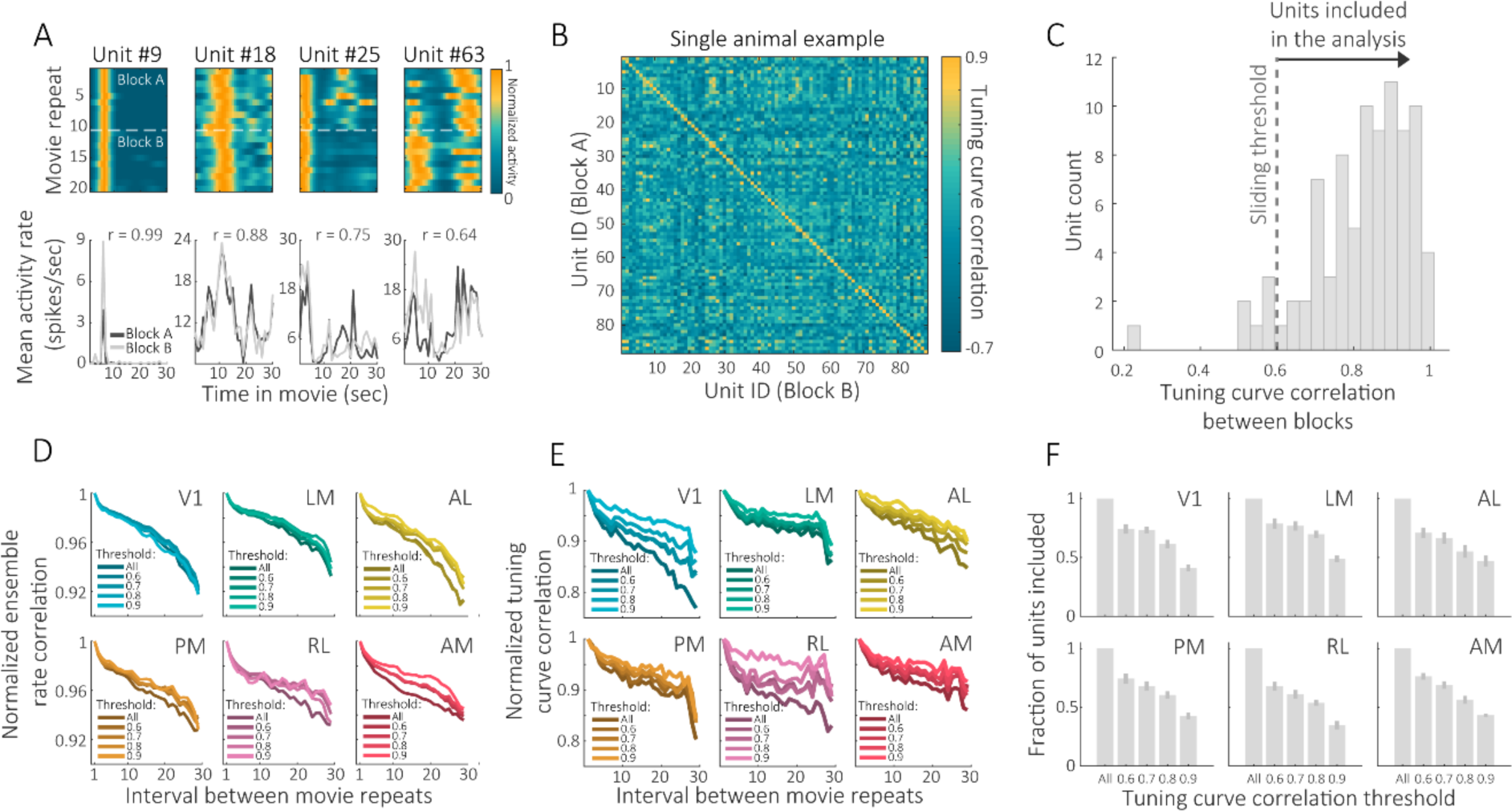
Representational drift is not a result of recording instability. **A**. Responsiveness of four V1 example cells from the same representative mouse across different repeats of ‘Natural movie 1’, spanning two blocks within the same recording session. Each unit exhibits a different degree of tuning curve stability across the two blocks (indicated by the Pearson correlation values in the bottom panels). **B**. Tuning curve correlation between blocks for all the units of the same representative mouse shown in A. **C**. Distribution of the tuning curve correlation values of the main diagonal in B. Units that show high tuning curve correlation across blocks are unlikely to represent cells whose identity is unstable within blocks. A sliding threshold was used to include different subsets of units with high tuning stability between blocks. **D**. Repeating the within-block stability analysis (shown in Fig. 2H) while subsampling units based on their tuning curve correlation between blocks. **E**. Repeating the within-block stability analysis (shown in Fig. 2I) while subsampling units based on their tuning curve correlation between blocks. **F**. Fraction of units included in the analysis as a function of their tuning curve correlation between blocks (mean ± SEM across mice from the Neuropixels ‘Functional connectivity’ group).

**Supplementary Figure 5.**
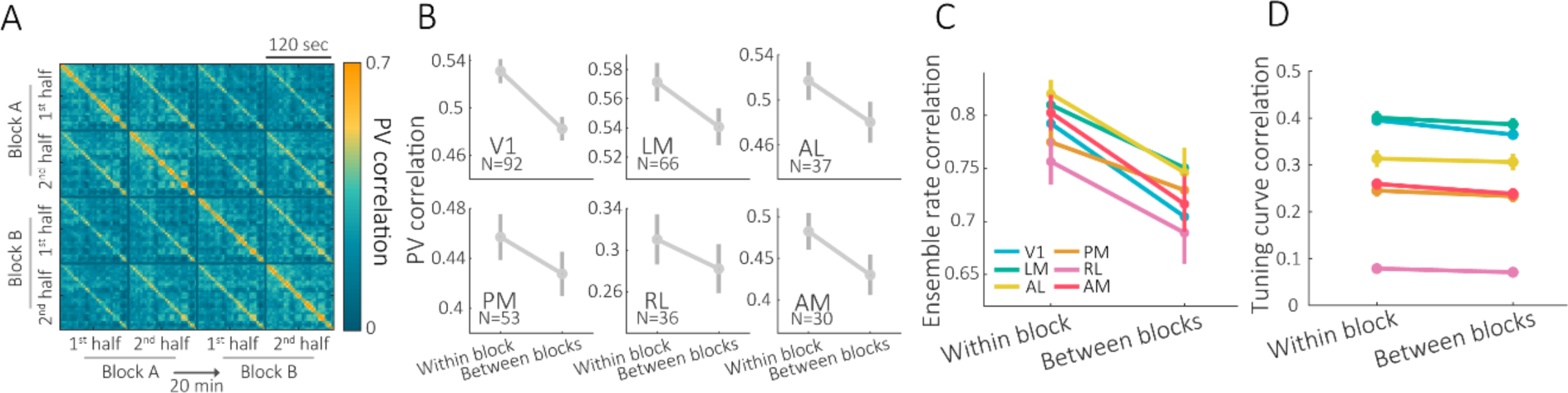
Visual representations change over timescales of tens of minutes. **A**. PV correlation between the 1^st^ (repeats 1-2) and 2^nd^ (repeats 3-5) halves of two different blocks of ‘Natural movie 3’ in a single visual area. The presented examples are the mean matrices across mice recorded in area AM using two-photon Ca^2+^ imaging. **B**. PV correlation between the two halves of the same block (within block) and between halves of different blocks (between blocks) using the Ca^2+^ imaging dataset (p < 10^−3^ for all areas, two-tailed Wilcoxon signed-rank test with Holm–Bonferroni correction). **C**. Ensemble rate correlation between the two halves of the same block (within block) and between halves of different blocks (between blocks) using the Ca^2+^ imaging dataset (p < 10^−3^ for all areas, two-tailed Wilcoxon signed-rank test with Holm–Bonferroni correction). **D**. Tuning curve correlation between the two halves of the same block (within block) and between halves of different blocks (between blocks) using the Ca^2+^ imaging dataset (p ≤ 0.03 for all areas, except areas AL and RL in which p > 0.05, two-tailed Wilcoxon signed-rank test with Holm–Bonferroni correction). Data in all panels are mean ± SEM across mice.

**Supplementary Figure 6.**
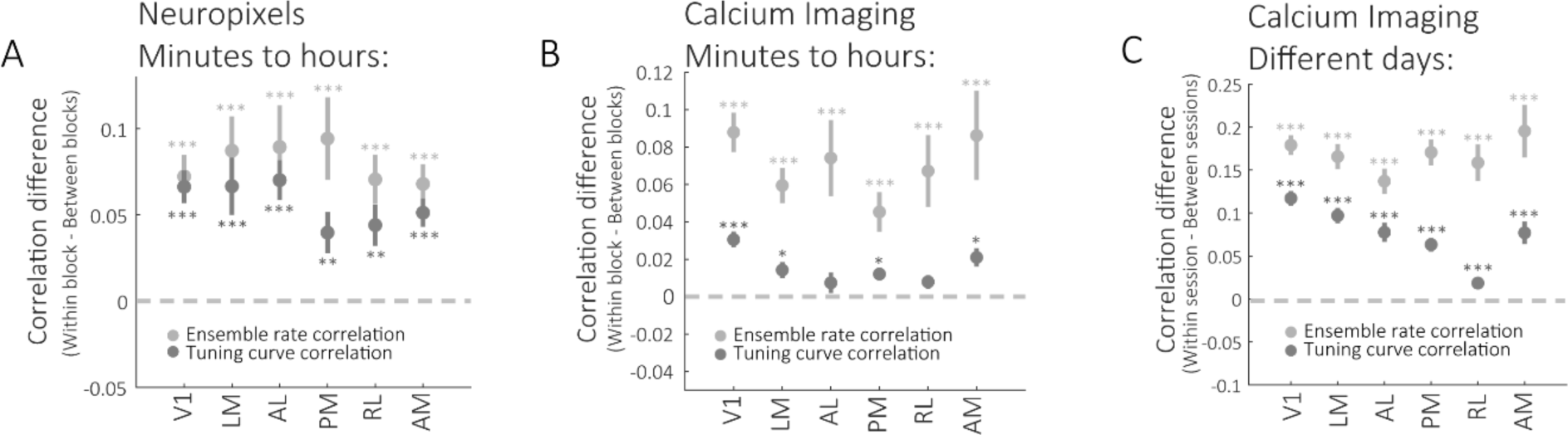
Magnitude of change in ensemble rate and tuning curve correlation as a function of time. **A**. The difference in ensemble rate and tuning curve correlations between ‘within block’ and ‘between blocks’ of ‘Natural movie 3’ for the Neuropixels dataset. **B**. The difference in ensemble rate and tuning curve correlations between ‘within block’ and ‘between blocks’ of ‘Natural movie 3’ for the Ca^2+^ imaging dataset. **C**. The difference in ensemble rate and tuning curve correlations between ‘within session’ and ‘between sessions’ of ‘Natural movie 1’ for the Ca^2+^ imaging dataset. Data are mean ± SEM across mice; two-tailed Wilcoxon signed-rank test with Holm–Bonferroni correction; * p<0.05, ** p<0.01, *** p<0.001, two-tailed.

**Supplementary Figure 7.**
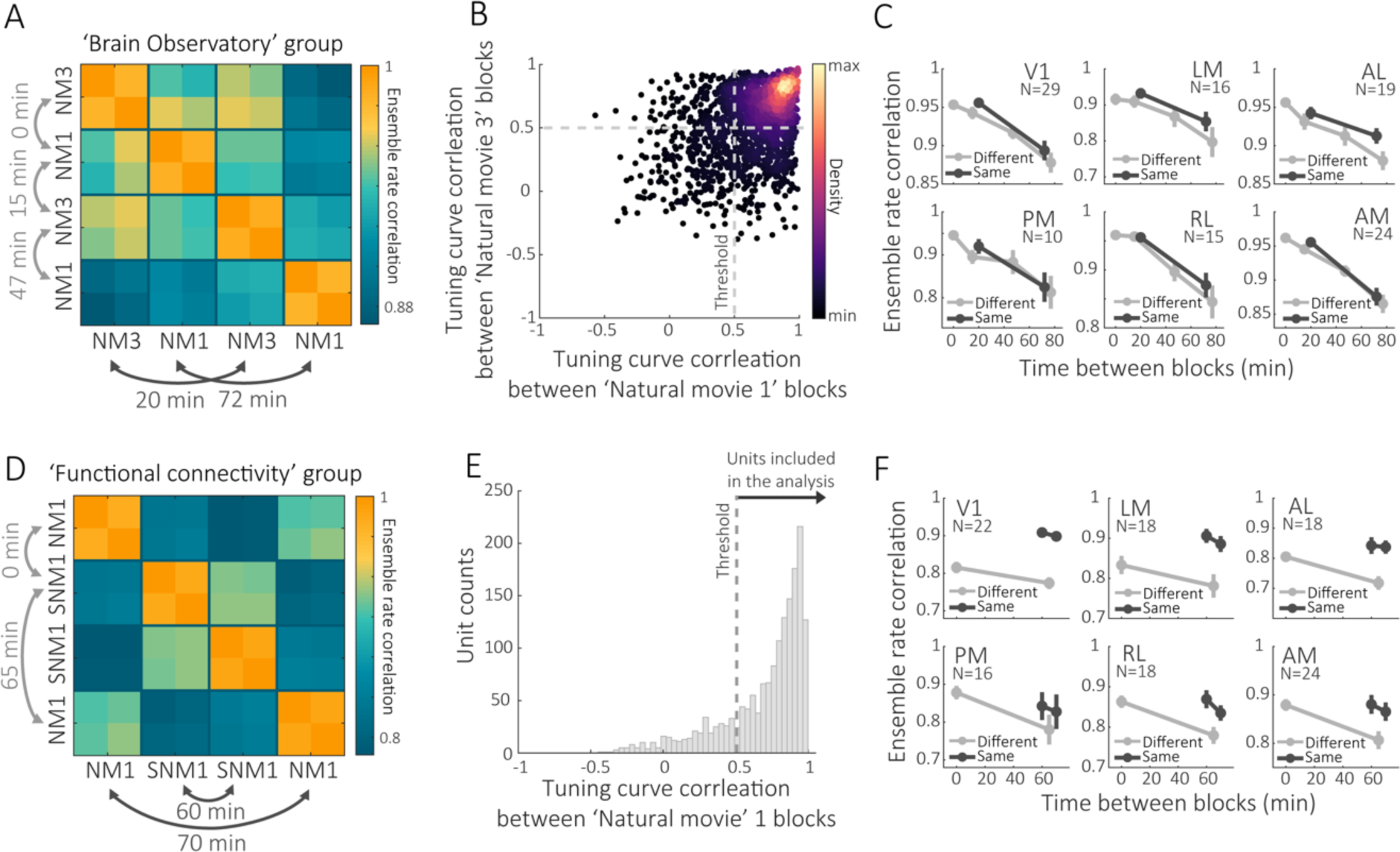
Ensemble rate correlations continuously decline over more than 1 hour. **A**. Ensemble rate correlations between halves of the same and different blocks within the same session (NM1, natural movie 1; NM3, natural movie 3). The presented example is the mean matrices across mice recorded with Neuropixels probes in area V1. **B**. Tuning curve correlations between blocks of ‘Natural movie 1’ and ‘Natural movie 3’ for all V1 units across mice. Each data point represents a single unit. The units included in the analysis are those with tuning curve correlation r>0.5 for both movies. **C**. Correlations between cell activity patterns across blocks of both the same and different blocks of natural movies decay with time. Note that ensemble rate correlations continuously decline with time, both between blocks of the same movie (dark gray) and between blocks of different movies (light gray). **D**. Ensemble rate correlations between halves of natural movie 1 and shuffled movie 1 blocks within the same session (NM1, natural movie 1; SNM1, shuffled natural movie 1). The presented example is the mean matrices across mice recorded with Neuropixels probes in area V1. **E**. The V1 units included in this analysis showed tuning curve correlation r>0.5 across the two blocks of ‘Natural movie 1’. **F**. Similarly, the ensemble rate correlations across different blocks of ‘Natural movie 1’ and different blocks of shuffled movie 1 declined with time.

**Supplementary Figure 8.**
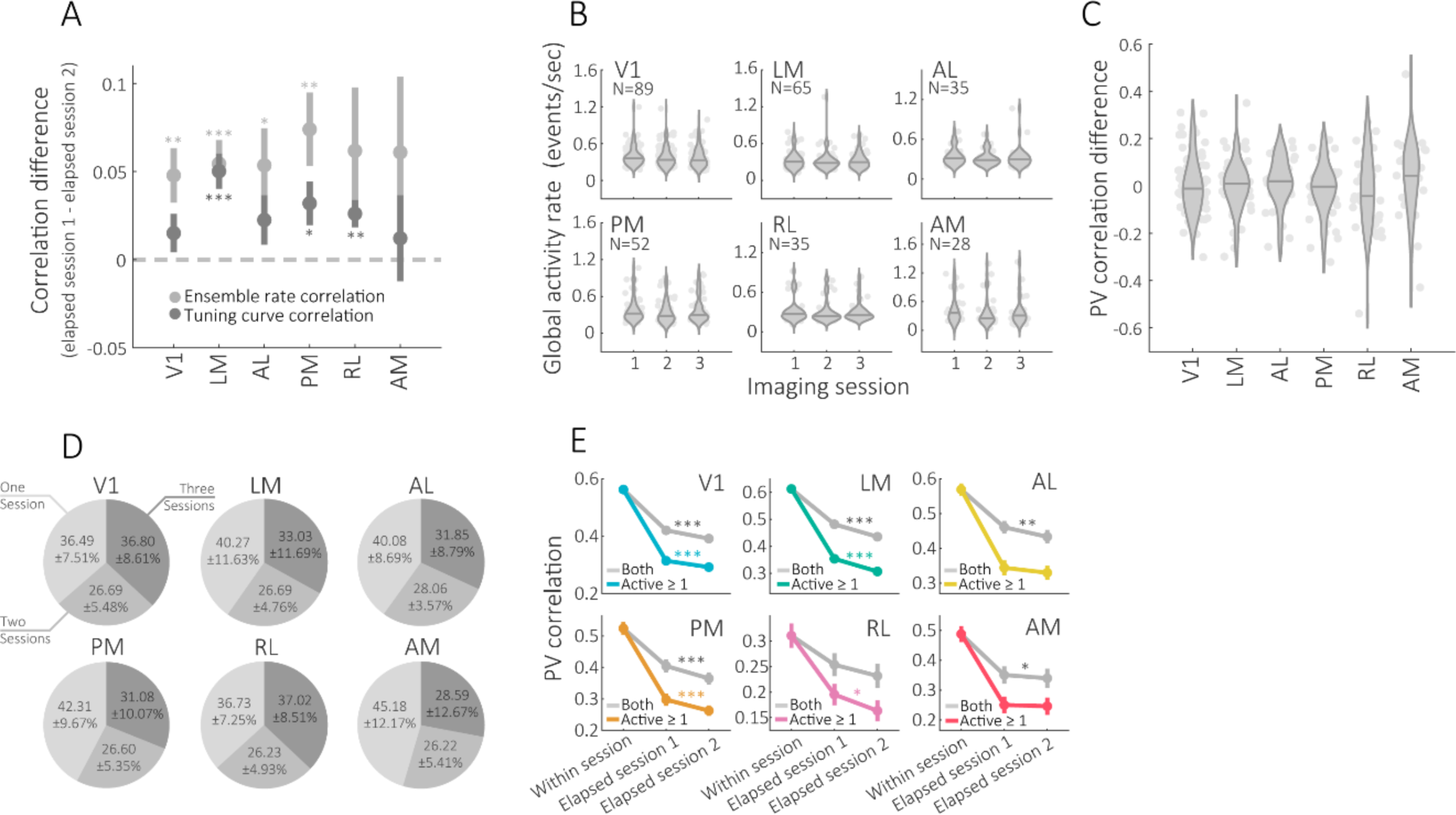
Stability of visual representations over days. **A**. The difference between ensemble rate and tuning curve correlations of temporally proximal sessions (the average correlation between sessions 1&2 and between sessions 2&3) and that of two distal sessions (the correlation between session 1&3); V1 (Z_Rate_ = 3.31, p = 0.002; Z_Tuning_ = 2.09, p = 0.053), LM (Z_Rate_ = 4.27, p < 10^−4^; Z_Tuning_ = 4.39, p < 10^−4^), AL (Z_Rate_ = 2.57, p = 0.014; Z_Tuning_ = 1.77, p = 0.075), PM(Z_rate_ = 3.34, p = 0.002; Z_tuning_ = 2.65, p = 0.0159), RL (Z_rate_ = 1.53, p = 0.068; Z_tuning_ = 3.03, p = 0.005), AM (Z_rate_ = 1.83, p = 0.068; Z_tuning_ = 0.87, p = 0.19), one-tailed Wilcoxon signed-rank test with Holm–Bonferroni correction; *p < 0.5, ** p < 0.01; ***p < 0.001. **B**. Distribution of population mean activity rates (Ca^2+^ events/frame) across mice for each of the imaging sessions. **C**. Distribution of the differences in the PV correlation values between pairs of subsequent sessions (i.e., the similarity between sessions 1 and 2 compared to that of sessions 2 and 3). Each data point represents an individual mouse; V1 (t_88_ = 0.07, p = 0.52), LM (t_64_ = −0.06, p = 0.47), AL (t_34_ = 0.87, p = 0.8), PM (t_51_ = −0.83, p = 0.2), RL (t_34_ = −0.77, p=0.22), AM (t_27_ = 1.21, p=0.86), one-tailed paired *t* test without correction for multiple comparisons. **D**. Percentage of cells active in a one, two or all imaging sessions. Mean ± SD across mice. **E**. Repeating the analysis presented in Fig. 3I for cells active in both compared time points (‘active both’), and for cells that were active in at least one of the compared time points (‘active ≥ 1’); post-hoc one-tailed Wilcoxon signed-rank test for the difference between the value of elapsed session 1 and that of elapsed session 2; *p < 0.5, ** p < 0.01; ***p < 0.001. Data in panels A and E are mean ± SEM across mice.

**Supplementary Figure 9.**
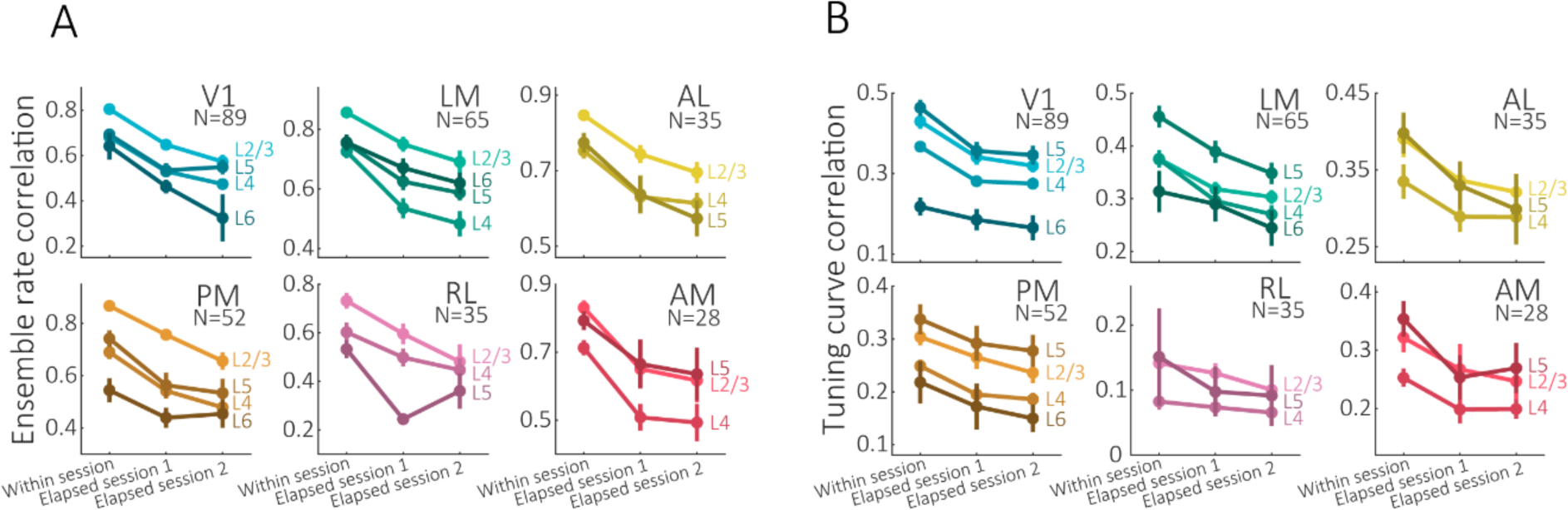
Comparison of representational drift between different cortical areas and layers across days. **A**. Ensemble rate correlation between the two halves of the same session (within session), between halves of two temporally proximal sessions (elapsed session 1) and between halves of two temporally distal sessions (elapsed session 2). **B**. Tuning curve correlation between the two halves of the same session (within session), between halves of two temporally proximal sessions (elapsed session 1) and between halves of two temporally distal sessions (elapsed session 2). Colors indicate different cortical layers. Data are mean ± SEM across mice.

**Supplementary Figure 10.**
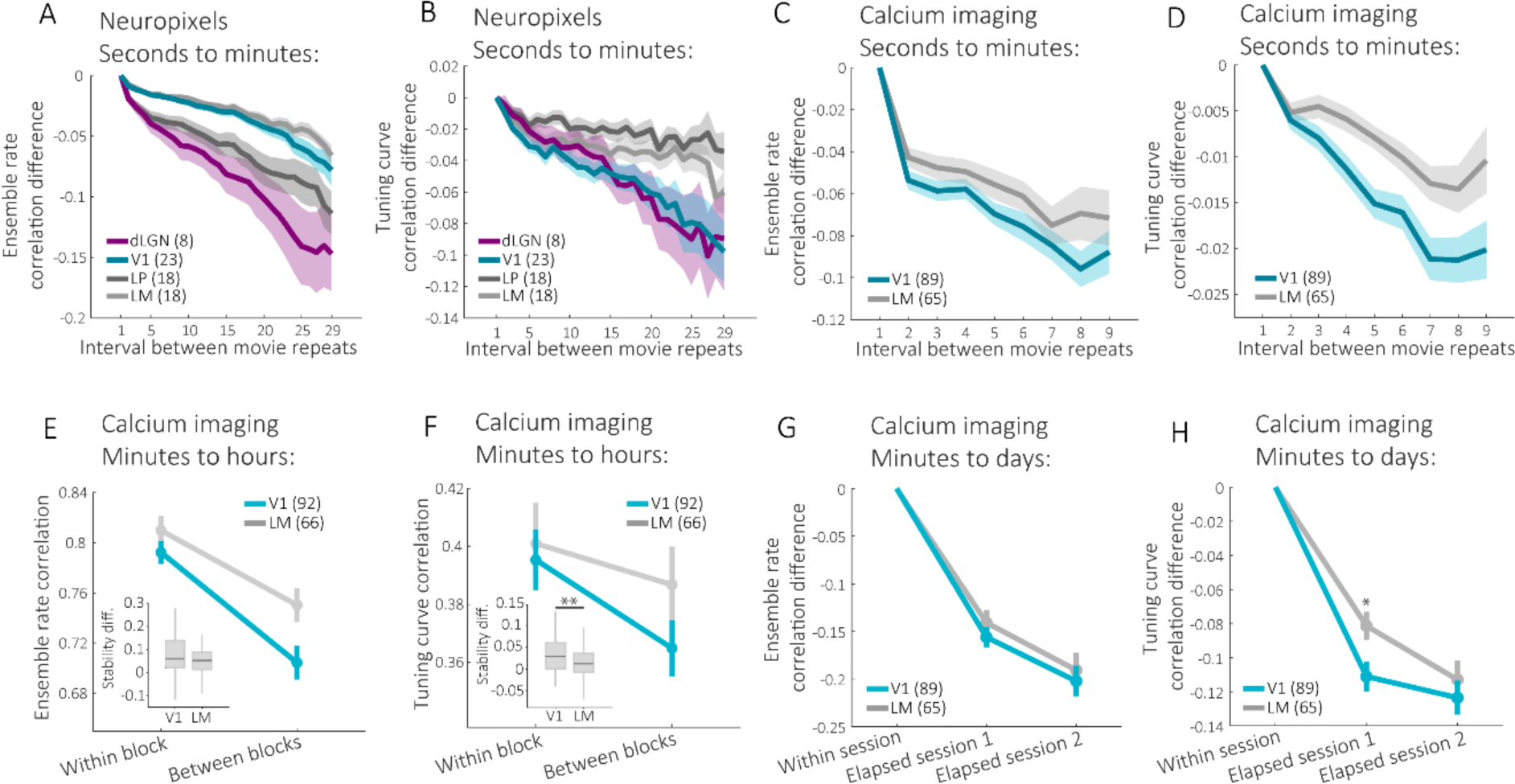
Comparison of representational drift between lower and higher visual areas. **A**. Difference in the ensemble rate correlation as a function of the interval between movie repeats for dLGN, LP, V1 and LM; for V1-LM there was a significant main effect for the interval between movie repeats (F_(1.9, 74.41)_ = 12.24, p < 10^−4^) and a non-significant interaction between area and the interval between movie repeats (F_(1.9, 74.41)_ = 1.244, p = 0.29); for LGN-LP there was a significant main effect for the interval between movie repeats (F_(1.82, 43.78)_ = 6.86, p = 0.003) and a non-significant interaction between area and the interval between movie repeats (F_(1.82, 43.78)_ = 1.63, p = 0.2), two-way mixed model ANOVA with Greenhouse-Geisser correction. **B**. Difference in the tuning curve correlation as a function of the interval between movie repeats for the same areas shown in A; for V1-LM there was a significant main effect for the interval between movie repeats (F_(2.12, 82.92)_ = 6.48, p = 0.0019) and a non-significant interaction for area and the interval between movie repeats (F_(2.12, 82.92)_ = 1.98, p = 0.14), for LGN-LP there was a significant main effect for the interval between movie repeats (F_(3,72.13)_ = 9.13, p < 10^−4^) and a significant interaction between area and the interval between movie repeats (F_(3, 72.13)_ = 5.32, p = 0.002), two-way mixed model ANOVA with Greenhouse-Geisser correction. **C**. Difference in the ensemble rate correlation as a function of the interval between movie repeats; there was a significant main effect for the interval between movie repeats (F_(3.32, 504.64)_ = 9.04, p < 10^−5^) and a non-significant interaction between area and the interval between movie repeats (F_(3.32, 504.64)_ = 0.89, p = 0.45), two-way mixed model ANOVA with Greenhouse-Geisser correction. **D**. Difference in the tuning curve correlation as a function of the interval between movie repeats; there was a significant main effect for the interval between movie repeats (F_(3.25, 495.47)_ = 11.48, p < 10^−4^) and a significant interaction between area and the interval between movie repeats (F_(3.25, 495.47)_ = 5.32, p = 0.047), two-way mixed model ANOVA with Greenhouse-Geisser correction. **E**. Ensemble rate correlation between the two halves of the same block (‘within block’) and between halves of different blocks (‘between blocks’). Inset: the difference in ensemble rate correlation between ‘within block’ and ‘between blocks’ for V1 compared to LM; two-tailed Mann–Whitney rank-sum test, Z = 1.63, p = 0.101. **F**. Tuning curve correlation between the two halves of the same block (‘within block’) and between halves of different blocks (‘between blocks’). Inset: the difference in tuning curve correlation between ‘within block’ and ‘between blocks’ for V1 compared to LM; two-tailed Mann–Whitney rank-sum test, Z = 2.6, p = 0.009. **G**. Ensemble rate correlation between the two halves of the same session (‘within session’), between halves of two temporally proximal sessions (‘elapsed session 1’) and between halves of two temporally distal sessions (‘elapsed session 2’). **H**. Tuning curve correlation between the two halves of the same session (‘within session’), between halves of two temporally proximal sessions (‘elapsed session 1’) and between halves of two temporally distal sessions (‘elapsed session 2’); the difference in tuning curve correlation between ‘within session’ and ‘between sessions’ for V1 compared to LM; ‘Within session’-‘Elapsed session 1’ (Z = −2.3, p = 0.042), ‘Within session’-‘Elapsed session 2’ (Z = −0.95, p = 0.34); two-tailed Mann– Whitney rank-sum tests with Bonferroni correction. Data in all panels are mean ± SEM across mice. The number of mice recorded from each area is indicated in parentheses. In A-D and G-H, correlations were scaled by subtracting the correlation value at the minimal interval between movie presentations.

**Supplementary Figure 11.**
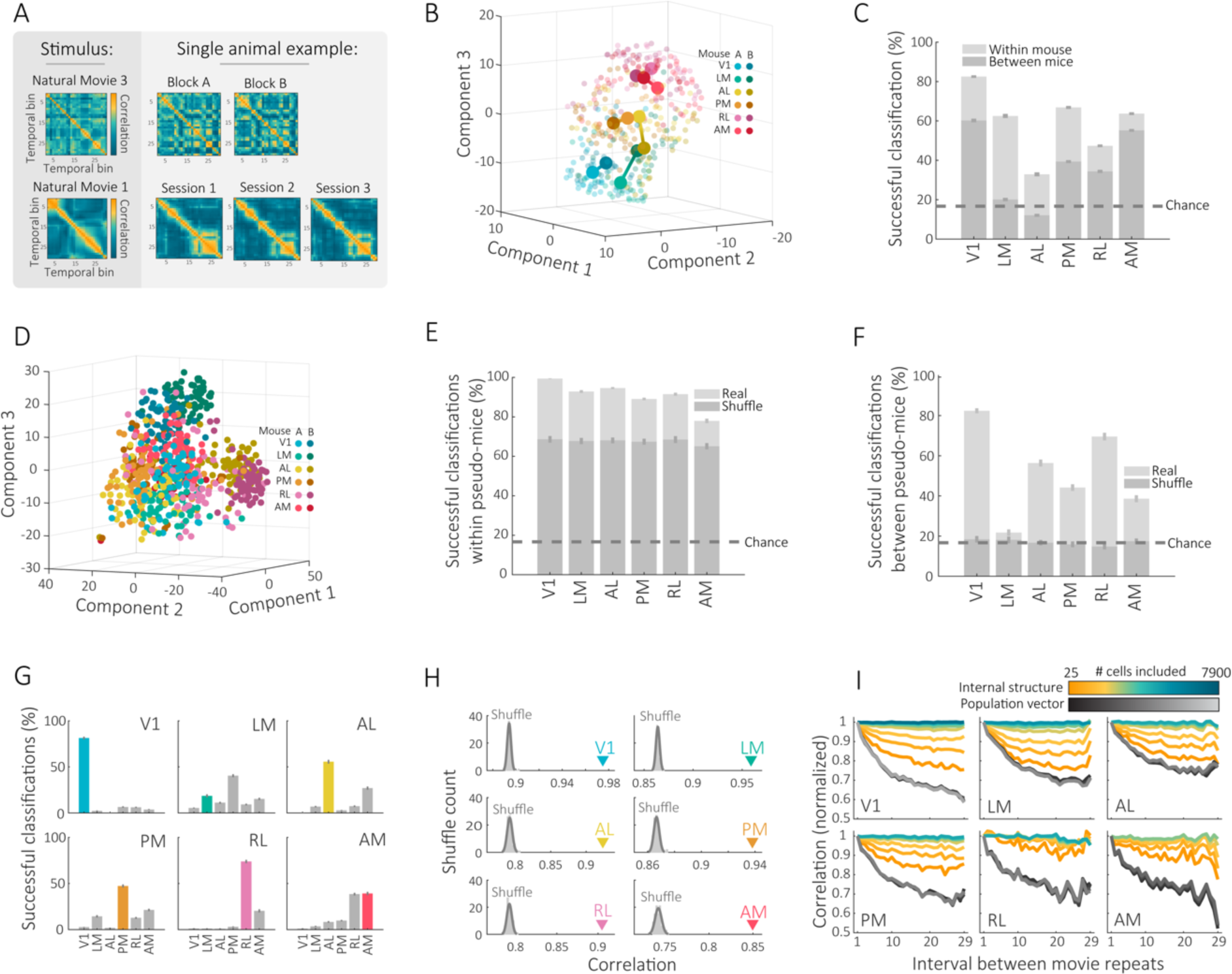
The unique coding properties of each visual area underlie a stereotypic internal structure of neuronal population activity. **A**. Different natural movies have a distinct structure of correlations between their individual frames (binned into 30 equally spaced segments). Each movie elicited a different structure of correlations between the population activities at each temporal bin (i.e., an internal structure of neuronal population activity). This structure of similarities between neuronal representations appears to be stable across different blocks within a session (top), and between different sessions across multiple days (bottom). Data is from a single mouse recorded using two-photon Ca^2+^ imaging in area LM. **B**. Example for dimensionality reduction (tSNE) on the internal structures of ‘Natural movie 1’ produced from two Ca^+2^ imaging ‘pseudo-mice’. Each data point corresponds to an internal structure of a single ‘Natural movie 1’ repeat. **C**. Percentage of successful classifications of the internal structures to their corresponding visual areas within and across Ca^+2^ imaging pseudo-mice (mean ± SEM across n=100 pairs of pseudo-mice). **D**. Example for dimensionality reduction (tSNE) on the internal structures of ‘Natural movie 1’ produced from two ‘shuffled pseudo-mice’ (mice A & B), which were generated from the same ‘pseudo-mice’ presented in Fig. 4C. Each data point corresponds to a single ‘Natural movie 1’ repeat. **E**. Percentage of successful classifications of internal structures across brain areas of the same ‘pseudo-mice’ compared to ‘shuffled pseudo-mice’ and chance. Internal structures from all areas exhibited significantly higher success in classification compared to their corresponding areas in the shuffled mice and chance (n=100 pairs of pseudo-mice; p ≤ 0.0033 for all areas, two-tailed Wilcoxon signed-rank test with Holm–Bonferroni correction). **F**. Percentage of successful classifications of internal structures between brain areas of different ‘pseudo-mice’ compared to ‘shuffled pseudo-mice’ and chance. All areas except LM exhibited significantly higher success in classification between mice compared to their corresponding areas in the shuffle mice and chance (n=100 pairs of pseudo-mice; p < 10^−9^ for all areas, two-tailed Wilcoxon signed-rank test with Holm–Bonferroni correction). **G**. Distribution of internal structure classifications between brain areas of different ‘pseudo-mice’. **H**. Mean internal structure stability compared to the distribution of temporally shuffled internal structures across areas from pseudo-mice created using the calcium imaging dataset. **I**. Correlation between the internal structures (colored lines) or the PVs (gray lines) as a function of the distance between repeats of ‘Natural movie 1’, plotted for varying numbers of neurons included in the analysis. Correlations were normalized to the value at the minimal interval between movie repeats.

**Supplementary Figure 12.**
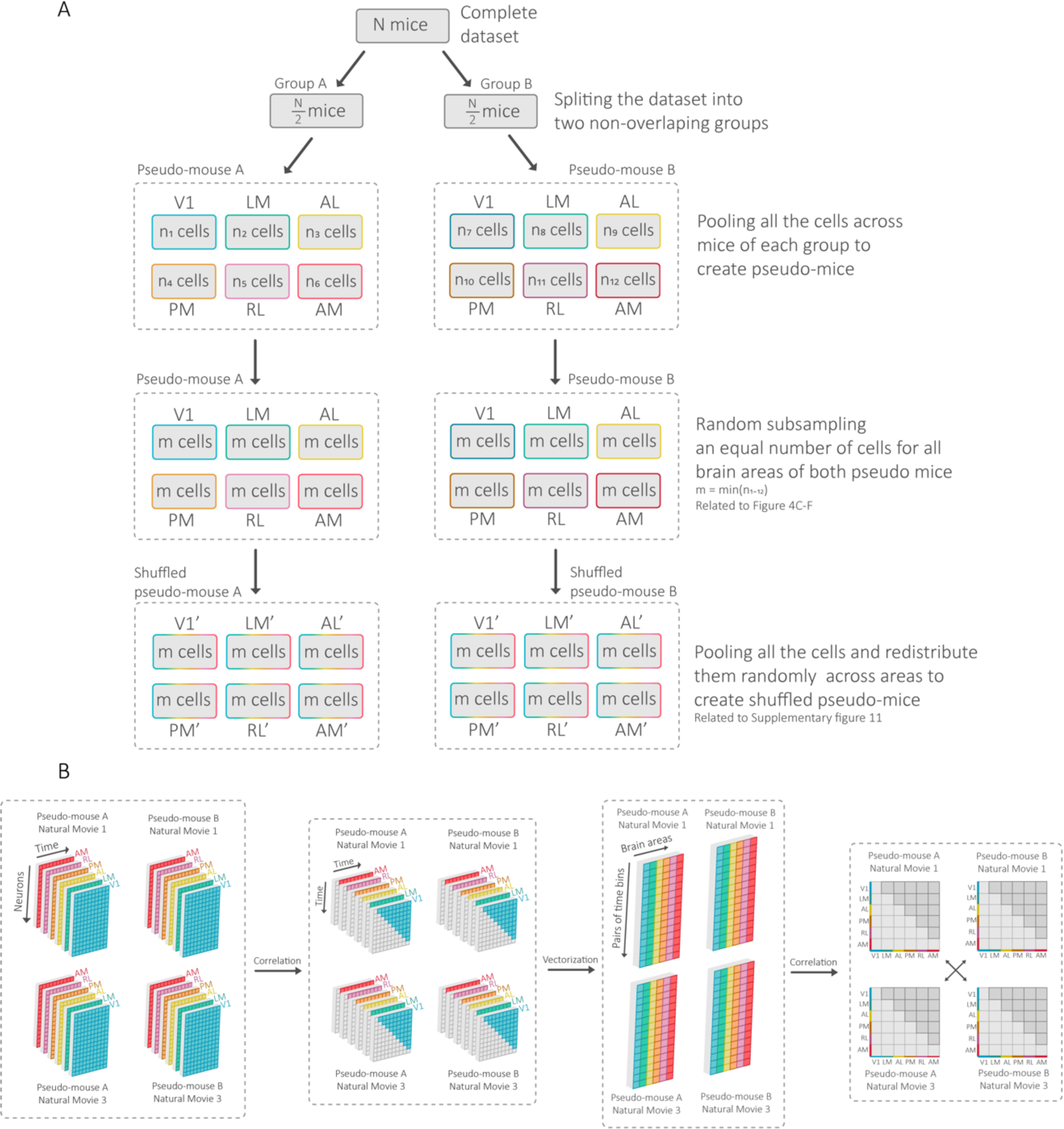
Generation of pseudo-mice and shuffled (control) pseudo-mice. **A**. Schematic workflow of the generation of the ‘pseudo-mice’ and ‘shuffled pseudo-mice’ presented in Fig. 4 and Supp. Fig. 11, respectively. We randomly divided a complete dataset of N mice into two equal, non-overlapping, groups of N/2 mice. Each mouse contained the neuronal activity recorded from 1-6 brain areas (depending on the dataset). Pooling all the cells related to each visual area across all mice yielded six ‘pseudo-areas’ for each of the ‘pseudo-mice’ (twelve ‘pseudo-areas’ in total). Because of variability in the number of recorded areas and cells across mice in the original dataset, the pooling procedure can result in a different number of cells in each of the ‘pseudo-areas’. To eliminate the possibility that differences between the internal structure of neuronal activity across visual areas stem from sampling biases, each ‘pseudo-area’ was randomly subsampled to contain an equal number of cells. The exact number of sampled cells was determined based on the area with the lowest cell count across ‘pseudo-mice’. Lastly, a complementary ‘shuffled pseudo-mouse’ was generated by the random redistribution of all the cells across ‘pseudo-areas’ in each of the ‘pseudo-mice’. This step yielded ‘shuffled pseudo-areas’ that served to evaluate the unique contribution of each area’s coding properties to the relationships within and between visual areas. **B**. Schematic workflow of the analysis performed to measure the degree to which the relationship between the internal structures of different areas is conserved across different movies and pseudo-mice (shown in Fig. 4E). Starting with 24 equally sized matrices (n x t) containing the mean neuronal activity across movie repeats in each temporal bin for each brain area, natural movie and pseudo-mouse (6 areas x 2 natural movies x 2 pseudo-mice). Correlating each temporal bin with the rest of the bins within a matrix produces equally sized (t x t) matrices for each brain area. Vectorizing the upper half of these matrices produces vectors representing the internal structures (vector size = (t^2^-t)/2)). Then, for each natural movie within a given pseudo-mouse, we calculated the Pearson’s correlation matrix across the internal structures of all areas. This procedure yielded four matrices (2 natural movies x 2 pseudo-mice), each symmetric and 6-by-6 in size (across all areas). Finally, we calculated the Pearson’s correlation between the matrices (using the vectorization of the upper half of each matrix) of different natural movies and different pseudo-mice, and averaged across the two comparisons (correlation between ‘Natural movie 1’ in pseudo-mouse A with that of ‘Natural movie 3’ in pseudo-mouse B, and vice versa).

